# The MICOS Complex Regulates Mitochondrial Structure and Oxidative Stress During Age-Dependent Structural Deficits in the Kidney

**DOI:** 10.1101/2024.06.09.598108

**Authors:** Prasanna Katti, Praveena Prasad, Sepiso K. Masenga, Prasanna Venkhatesh, Zer Vue, Andrea G. Marshall, Benjamin Rodriguez, Han Le, Edgar Garza-Lopez, Alexandria Murphy, Brenita Jenkins, Ashlesha Kadam, Jianqiang Shao, Amber Crabtree, Pamela Martin, Chantell Evans, Mark A. Phillips, David Hubert, Nelson Wandira, Okwute M. Ochayi, Dhanendra Tomar, Clintoria R. Williams, Jennifer Gaddy, Briar Tomeau, LaCara Bell, Taneisha Gillyard, Markis’ Hamilton, Vineeta Sharma, Mohd Mabood Khan, Elma Zaganjor, Olujimi A. Ajijola, Estevão Scudese, Tyne W. Miller Fleming, André Kinder, Chandravanu Dash, Anita M. Quintana, Bret C. Mobley, Julia D. Berry, Pooja Jadiya, Dao-Fu Dai, Annet Kirabo, Oleg Kovtun, Jenny C. Schafer, Sean Schaffer, Renata Oliveira Pereira, Melanie R. McReynolds, Antentor Hinton

## Abstract

Due to aging, the efficiency of kidney function begins to decrease. Dysfunction in mitochondria and their cristae is a hallmark of aging. Therefore, age-related decline in kidney function could be attributed to changes in mitochondrial ultrastructure, increased reactive oxygen species, and alterations in metabolism and lipid composition. We sought to understand how mitochondrial ultrastructure is altered over time in tubular kidney cells. A serial block facing-scanning electron microscope and manual segmentation using the Amira software were employed to visualize murine kidney samples during the aging process at 3 months (young) and 2 years (old). We found that 2-year mitochondria are more fragmented with many uniquely shaped mitochondria observed across aging, concomitant with shifts in ROS, metabolomics, and lipid homeostasis. Furthermore, we demonstrate that the mitochondrial contact site and cristae organizing system (MICOS) complex is impaired in the kidney during aging. Disruption of the MICOS complex resulted in altered mitochondrial metabolic function and increased ROS levels. We found significant, detrimental structural changes in the mitochondria of aged kidney tubules, suggesting a potential mechanism underlying the increased frequency of kidney disease with aging. We hypothesize that disruption of the MICOS complex exacerbates mitochondrial dysfunction, creating a vicious cycle of mitochondrial degradation and oxidative stress, which impacts kidney health.

**Impact and Implications:** Due to aging, the efficiency of kidney function begins to decrease, and the risk of kidney diseases may increase; however, the specific regulators of mitochondrial age-related changes are poorly understood. This study demonstrates that the MICOS complex may be a target for mitigating age-related mitochondrial changes. The MICOS complex is associated with oxidative stress and calcium dysregulation, which also arise in many kidney pathologies.

**Highlights:**

- Aging alters the MICOS mRNA levels and disease markers.
- Aging reduces cristae architecture, mitochondrial volume and complexity in murine kidney ultrastructure
- Reducing MIC60 and CHCHD6 lowers Ca^2+^ uptake and retention and induces oxidative stress in HEK cells.
- Metabolomic Profiling revealed that NAD^+^ and amino acid metabolism were altered in aged kidneys.
- MICOS deficiency alters the reduced basal, ATP-linked, maximal capacity and spare capacity.
- Decreased modeled expression of *CHCHD6* in individuals of European genetic ancestry is linked to chronic kidney disease, whereas decreased modeled expression of *OPA1* in individuals of African genetic ancestry is associated with chronic kidney disease.

**Graphical Abstract:** Kidney aging causes a decline in the MICOS complex, concomitant with metabolic, lipidomic, and mitochondrial structural alterations.

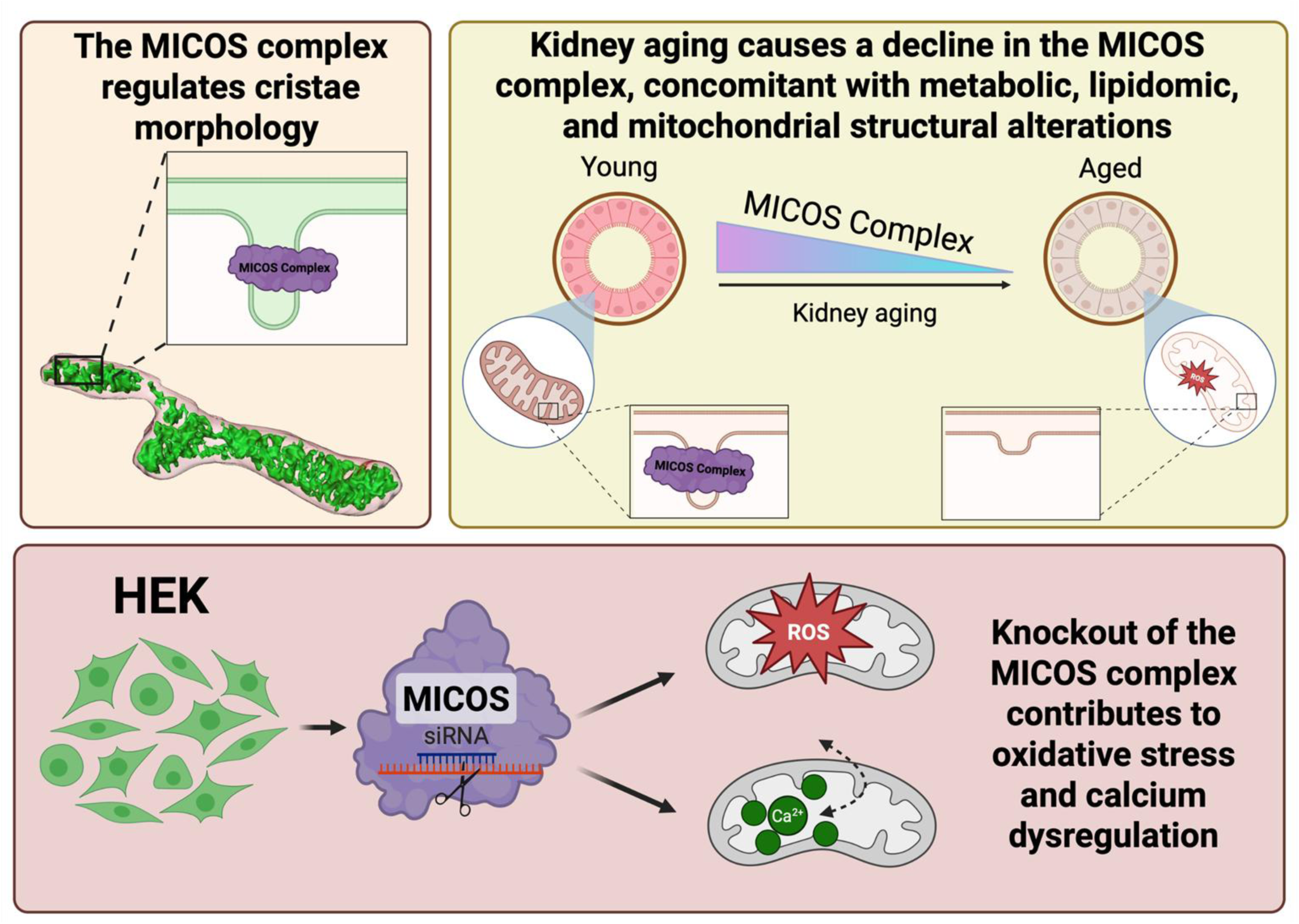

## INTRODUCTION

Kidneys are primarily known for their role in excreting waste products from the body. However, their functions extend far beyond this, including hormonal signaling, making the kidney critical for many other functions, such as blood pressure regulation^1^. However, renal dysfunction may occur in various states, such as the sudden loss of kidney function, acute kidney injury (AKI), and chronic kidney diseases (CKD)^2^. An estimated 90% of afflicted individuals are not aware that they have an AKI or CKD; thus, the exact prevalence and burden of kidney diseases remain difficult to measure^3^. Poor treatment outcomes for kidney diseases and their associations with other conditions, including cardiovascular disease, emphasize the importance of developing new, effective treatments for kidney disease^4^. Kidneys are among the most mitochondria-rich tissues in the body^5^. Therefore, studying the role of mitochondria in kidney function and disease provides an avenue for understanding kidney disease pathophysiology.^5^

Researchers widely recognize that mitochondria provide numerous critical cellular functions beyond just energy production through oxidative phosphorylation^6^. Mitochondria play a crucial role in cell signaling, calcium regulation, apoptosis, and overall cellular homeostasis. Mitochondrial genes also encode the pathways responsible for ATP generation^7^. Mitochondrial dysfunction contributes to kidney diseases, ^8–11^ and is also implicated in diseases affecting mitochondria-rich organs, including muscle diseases ^12,13^, neurological diseases ^14,15^, and obesity-associated diabetes ^16,17^. Notably, specific mechanisms of mitochondrial dysfunction that may govern each disease state, as well as ways to rescue mitochondrial function, remain nascent research topics. Therefore, it is essential to investigate the therapeutic implications of mitochondria.

Studies have shown that mitochondrial dysfunction is a hallmark of AKI pathogenesis, making mitochondria a critical target for restoring kidney function to pre-disease states ^2,18^. Other focus areas for kidney research include autosomal dominant polycystic kidney disease, which has been shown to shift mitochondrial function toward anaerobic respiration^19^ through various mechanisms, such as calcium signaling, thereby limiting oxidative capacity. The primary roles, frequency, and connections of mitochondria in different tissues may vary significantly. For mitochondrial calcium regulation, the mitochondrial calcium uniporter (MCU), which regulates calcium influx in mitochondria, is more active in the kidney than in other mitochondria-rich organs, such as the liver and the heart ^20^. Still, the mechanisms that mediate kidney mitochondrial dynamics remain unclear.

One link between mitochondrial function and kidney diseases is the aging process. Aging is the most significant risk factor for both CKD and AKI^22–23^. It is well established that mitochondrial function declines with aging across organ systems^21–23^, with mitochondrial dysfunction now recognized as a hallmark of aging process^24^. Given the strong association between aging and the incidence of AKI and CKD, age-associated mitochondrial dysfunction is likely to contribute to the pathogenesis of both AKI and CKD ^21–23^. Past research has implicated that proton leak across the inner mitochondrial membrane leads to a loss of mitochondrial bioenergetics, reducing electron transfer chain efficiency in mouse kidneys^25^. Furthermore, clearance of damaged mitochondria through mitophagy responses is also blunted in aging kidney proximal tubules.^26^ However, it is poorly understood by what mechanism mitochondrial function declines with age, as well as the relationship between mitochondrial dysfunction and mitochondrial structural changes in the kidneys. It is possible that, as in other tissues, mitochondrial fusion is decreased with age, leading to damaged mitochondria and the breakdown of cristae. This is supported by previous studies examining aging kidneys via transmission electron microscopy (TEM) ^27^. However, TEM only allows analysis of mitochondria in 2D, which does not provide sufficient detail on mitochondrial subcellular structures^28,29^. Thus, we performed 3D reconstruction using a serial block face-scanning electron microscopy (SBF-SEM), which allows for a broader range of analysis,^30,31^ to determine how mitochondrial networking and structures are altered across aging in mouse kidneys. We studied mice at two ages: 3-month-old mice, representing a young adult phenotype, and 2-year-old mice, representing a geriatric model ^32^. We then studied the morphological changes in mitochondria and cristae in both 2D and 3D, as well as mitochondrial reactive oxygen species (ROS) production, in young and aged mice. Mitochondrial ROS synthesis mainly occurs on the electron transport chain located in the inner mitochondrial membrane during the process of oxidative phosphorylation^33,34^. Damaged or dysfunctional mitochondria are harmful to the cells because they release substances that promote cell death and create ROS, which causes Thymic epithelial cell-induced apoptosis^34^. The relationships among mitochondrial oxidative stress, ROS production, and mitophagy are closely intertwined, and all are involved in the pathophysiology of AKI ^34^.

Past studies have suggested that dysfunction of the MICOS complex, a protein complex that regulates cristae morphology, can lead to oxidative stress ^35^. Given that we have previously found age-dependent losses in the MICOS complex correlating with mitochondrial structural defects in both skeletal and cardiac muscle tissues, we hypothesized that a similar phenotype may be observed in the kidney^36,37^. The oxidative stress resulting from MICOS dysfunction during aging could confer oxidative damage and impaired calcium homeostasis, characteristic of age-related AKI and CKD^38,39^. We found that deletion of the MICOS genes resulted in mitochondrial structural changes and impairments in mitochondrial calcium regulation in the kidney. Because we demonstrated that aging affects both the structure of the kidneys and mitochondria, as well as MICOS, we investigated the metabolomic and lipidomic changes in aging kidneys to further understand the pathways that may mediate the roles of mitochondria and MICOS in aging kidney vulnerability to injury and disease.

## METHODS

### Animal Care and Maintenance

As per the protocols previously described, the care and maintenance of the male C57BL/6J mice conformed to the National Institute of Health’s guidelines for the use of laboratory animals^36,40,41^. The University of Iowa’s Institutional Animal Care and Use Committee (IACUC) or the University of Washington IACUC approved the housing and feeding of these mice. The mice were anesthetized using a mixture of 5% isoflurane and 95% oxygen. All young and old mice were sourced from the same institution and maintained on an identical standard chow diet composed of ground wheat, ground corn, ground oats and meat meals.

### MICOS complex GREX association with kidney disease phenotypes in BioVU

The Vanderbilt Institute for Clinical and Translational Research at Vanderbilt University Medical Center curates BioVU, a biobank of genotype and linked de-identified electronic health record (EHR) data. The BioVU program has been previously described^42,43^. Briefly, this opt-in program collects leftover blood samples during routine patient care visits at clinics across Tennessee and processes them for genotyping or genetic sequencing. BioVU currently houses samples from 344,467 individuals, with ongoing sample collection.

Genotype data for 94,474 individuals were generated on the Illumina Multi-Ethnic Genotyping Array (MEGAEX) as previously described.^44^ Quality control procedures included filtering for SNP and individual call rates, sex discrepancies, and excessive heterozygosity. We used principal component analysis to identify individuals of European and African genetic ancestry, using the 1000 Genomes populations as reference^45,46^. After imputation using the Michigan Imputation server and the Haplotype Reference Consortium (HRC) reference panel, genotype data underwent additional quality control procedures, including filtering for imputation quality, minor allele frequency, and Hardy-Weinberg Equilibrium^47,48^. After removing genetically related individuals, we identified 65,363 individuals of European genetic ancestry and 12,313 individuals of African genetic ancestry for analysis. We calculated the genetically regulated gene expression (GREX) in BioVU individuals using models built from the genotype-tissue expression (GTEx) version 8 project data, which includes genotype and matched transcriptome data for 838 donors across 49 distinct tissues^49^. The best-performing GREX models, based on the highest r^2^ values from PrediXcan, UTMOST, and JTI approaches, were used to model gene expression for *CHCHD3*, *CHCHD6*, and *OPA1* from BioVU genotype data^50–52^.

To evaluate correlations between kidney disease phenotypes and *CHCHD3*, *CHCHD6*, and *OPA1* GREX, we extracted kidney disease case status from de-identified EHR data in BioVU participants. Specifically, we tested phenotypes or phecodes mapped from ICD9/10 (International Classification of Diseases, 9th and 10th editions) billing codes (phenotypes tested: Renal failure NOS, CKD, and kidney replaced by transplant in all individuals and Renal sclerosis and Abnormal results of function study of kidney in individuals of European genetic ancestry only), requiring at least 50 cases for each phenotype analysis. We used previously described mapping of phecodes to ICD9/10 codes, which are available through the PheWAS package in R (version 0.99.5-2 and 3.6.0, respectively) and the online PheWAS catalogue^53,54^.

(https://phewascatalog.org/phewas/#home)

We required individuals to have at least two documented instances of the ICD codes on unique dates within the medical record to be considered a case. Controls had no relevant ICD codes in their medical records. Logistic regression analysis was used to examine the association between *CHCHD3*, *CHCHD6*, and *OPA1* GREX (predictor variable) with kidney phenotype case status (outcome variable). Covariates within the regression models included principal components (PC1-10), sex, current age, median age of the medical record, and genotype batch in the ancestry-stratified analyses. The most stringent global Bonferroni p-value adjustment was calculated by correcting the p-value for all gene-tissue GREX pairs tested (n=80) and all phenotypes tested (n=5, p=0.05/400). Because the GREX models across tissues are highly correlated, we also generated a less stringent within-tissue Bonferroni p-value adjustment, correcting the p-value for the number of genes tested (n=3) and the number of phenotypes tested (n=5 in European ancestry, p=0.05/15, and n=3 in African ancestry, p=0.05/9). Nominal significance was considered p<0.05.

### Human Sample Cohort

All human samples were obtained from Brazilian cohorts in accordance with the CARE (Ethics Appreciation Presentation Certificate) guidelines. Samples from young individuals were collected, and experiments were performed under CAEE number 61743916.9.0000.5281; samples from older individuals were collected under CAEE number 10429819.6.0000.5285.

### Immunohistochemistry

Young (4–5 months old) and old (21–23 months old) C57BL/6J were maintained on a chow diet. Kidney slices were embedded in OCT, with section processing as previously described^55^. Nitrotyrosine (MilliporeSigma, #06-284,1:1000), and anti-mouse IgG-HRP (Abcam; ab97046) staining were performed to measure oxidative stress. Masson trichrome staining was quantified using representative low-power images from each kidney section, deconvoluted and thresholded in ImageJ to calculate the blue area relative to the total tissue area.

### SBF-SEM Processing of Kidney Tissue

SBF-SEM was performed according to previously defined protocols ^56–58^. Anaesthesia was induced in male mice using 5% isoflurane. The extracted kidney tissue was treated with 2% glutaraldehyde in 100 mM phosphate buffer for 30 minutes, dissected into 1-mm³ cubes, and further fixed in a solution containing 2.5% glutaraldehyde, 1% paraformaldehyde, and 120 mM sodium cacodylate for 1 hour. Fixation and subsequent steps were collected onto formvar-coated slot grids (Pella, Redding, CA), stained and imaged as previously described ^56–58^. This includes tissue washing with 100 mM cacodylate buffer, incubation in a mixture of 3% potassium ferrocyanide and 2% osmium tetroxide, followed by dehydration in an ascending series of acetone concentrations. The tissues were then embedded in Epoxy Taab 812 hard resin. Sectioning and imaging of the sample were performed using a VolumeScope 2 SEM (Thermo Fisher Scientific, Waltham, MA).

### Live-Cell Imaging and Analysis

Live-cell mitochondrial dynamics were visualized using a Nikon Eclipse Ti2 inverted fluorescence microscope equipped with a Yokogawa CSU-W1 spinning disk confocal scanner, Hamamatsu Fusion BT camera, SoRa super-resolution module, environmental chamber, piezo stage controller, and solid-state lasers (405, 488, 561, and 640 nm), all controlled via NIS-Elements AR software (version 5.42). Cells cultured in 35-mm MatTek glass-bottom dishes were imaged with a 100× Plan Apo Lambda D oil immersion objective (NA 1.45), capturing MitoTracker Orange-labeled mitochondria through the 561 nm channel at 2.5-second intervals over 5 minutes using either standard W1 mode (xy pixel size: 65 nm) or SoRa mode (xy pixel size: 23 nm), with laser power maintained at 5% and 100 ms exposure time per frame to balance phototoxicity concerns with signal quality (SNR ∼1.5-2.5). The Perfect Focus System maintained z-stability throughout all acquisitions. At the same time, high-resolution z-stacks (10-20 µm thick) were acquired in SoRa mode with 100 nm step sizes and subsequently enhanced using the Nikon Batch Deconvolution module (version 6.10.02), which implements Blind and Richardson-Lucy algorithms with 20 iterations and automatic noise estimation.

### Assessment of ROS levels

HEK293 wild-type cells were cultured in high-glucose DMEM (4.5 g/L glucose) supplemented with 10% fetal bovine serum and 1% penicillin–streptomycin at 37 °C with 5% CO₂. Cells (0.2 × 10⁶) were plated in 35-mm dishes and transfected the following day with siRNAs targeting *MIC60* (Thermo Fisher, 136128) or *CHCHD6* (Thermo Fisher, 34035) using Lipofectamine RNAiMAX (Invitrogen), according to the manufacturer’s instructions. After incubation for 30 hrs., cells were co-stained for 30 minutes at 37°C with two different dyes for ROS detection: MitoBright ROS Deep Red (10 µM, Dojindo Laboratories) for mitochondrial superoxide detection, and DCFDA (10 µM, Invitrogen) for intracellular total ROS detection. Following incubation with staining dyes, cells were washed three times with 1X HBSS, and ROS imaging was performed using a confocal microscope (FV4000, Olympus Life Science).

For mitochondrial H_2_O_2_ imaging, cells were incubated with MitoPY1 (5 µM, Bio-Techne) for 45 min at 37°C. Cells were then washed with 1x HBSS and imaged using a confocal microscope (FV4000, Olympus Life Science). ImageJ was used for quantifying fluorescence intensities. COS7 cells were cultured in high-glucose DMEM (4.5 g/L glucose; Gibco) supplemented with 10% fetal bovine serum and 1% penicillin–streptomycin. Cells were treated with a MICOS inhibitor or vehicle control (DMSO) as indicated. Mitochondria were labelled using MitoTracker Orange according to the manufacturer’s instructions.

### Knockdown of *MIC60* and *CHCHD6* in HEK293 cells

The *MIC60* and *CHCHD6* siRNAs, along with a scramble siRNA control, were transfected into HEK293 cells using Lipofectamine RNAiMax (Invitrogen) according to the manufacturer’s instructions. After a 48-hour incubation, cells were used for Calcium measurements.

### Measurement of mitochondrial Ca^2+^ uptake in HEK293 cells

Mitochondrial Ca^2+^ uptake was assessed using a multi-wavelength excitation dual-wavelength emission fluorimeter (Delta RAM, Photon Technology Int.) with slight modifications following the protocol outlined in Tomar et al., 2016 (PMID: 27184846). An equal number of cells (2.5×10^6^ cells) were uniformly cleansed with Ca^2+^/Mg^2+^-free DPBS (GIBCO) and subsequently permeabilized in 1 mL of intracellular medium (ICM-120 mM KCl, 10 mM NaCl, 1 mM KH_2_PO_4_, 20 mM HEPES-Tris, pH 7.2) containing 20 μg/ml digitonin, 1.5 μM thapsigargin to inhibit the SERCA pump and 2.5 mM succinate to energize the mitochondria. The loading of Fura-FF (1 μM) at the 0s time point facilitated the measurement of mitochondrial Ca^2+^ uptake. Fluorescence was recorded at 340- and 380 nm excitation/510 nm emission, with continuous stirring at 37°C. At specified time points, a bolus of 5 μM Ca^2+^ and the mitochondrial uncoupler FCCP (10 μM) were introduced into the cell suspension.

### Assessment of mCa^2+^ retention capacity (CRC)

To assess mCa^2+^ retention capacity (CRC), 2 × 106 cells were resuspended in an intracellular-like medium containing (120 mM KCl, 10 mM NaCl, 1 mM KH2PO4, 20 mM HEPES-Tris, pH 7.2), 1.5 μM Thapsigargin (Tg) to inhibit SERCA so that the movement of Ca^2+^ was solely influenced by mitochondrial uptake, 20-μg/ml Digitonin (Dg), supplemented with 2.5 μM succinate. All solutions were treated with Chele× 100 (Sigma) to remove traces of Ca^2+^. Digitonin-permeabilized cells were loaded with the ratiometric reporter FuraFF at 1 μM. Fluorescence was recorded using a spectrofluorometer (Delta RAM, Photon Technology International) with excitation at 340 nm and emission at 510 nm. Following baseline recordings, a repetitive series of Ca^2+^ boluses (5 µM) was introduced at the indicated time points. Upon reaching a steady-state recording, a protonophore, 10 μM FCCP, was added to collapse the Δψm and release matrix-free Ca^2+^. The number of Ca^2+^ boluses taken up by cells was counted to calculate mitochondrial CRC.

### Data Analysis

GraphPad Prism (La Jolla, CA, USA) was used for all statistical analyses. All experiments involving SBF-SEM and TEM data included at least 3 independent replicates. Statistics were not handled by the experimenters. The black bars represent the standard error of the mean. For all analyses, one-way ANOVA was performed with tests against each independent group and significance was assessed using Fisher’s protected least significant difference (LSD) test. *, **, ***, **** were set to show significant difference, denoting *p* < 0.05, *p* < 0.01, *p* < 0.001, and *p* < 0.0001, respectively.

## RESULTS

### Genetic Association Analysis Reveals MICOS Complex Components as Potential Mediators of Kidney Disease Susceptibility

We conducted a comprehensive biobank-based investigation examining the relationship between genetically-determined expression profiles of MICOS complex components and clinical manifestations of kidney disease across a large patient cohort. Leveraging the GTEx v8 database containing matched genotype-transcriptome data from 838 donors across 49 tissues, researchers calculated genetically-regulated gene expression (GREX) for key mitochondrial structure regulators (*CHCHD3*, *CHCHD6*, and *OPA1*) in 77,676 BioVU participants (Figure 1A). Kidney disease cases and controls were identified using ICD9/10 diagnostic codes from de-identified electronic health records, followed by logistic regression analysis to test associations between GREX and kidney disease status while controlling for genetic ancestry (PC1-10), sex, age variables, and technical factors. The results demonstrate tissue-specific patterns of association, with *CHCHD6* GREX exhibiting significant relationships with multiple kidney phenotypes in European ancestry individuals (EUR), as visualized in the rank-ordered tissue-specific models (Figure 1B). Similarly, for African ancestry individuals (AFR), *OPA1* GREX showed notable associations with renal conditions, including chronic kidney disease, renal failure, kidney transplantation, and abnormal kidney function studies (Figure 1C), suggesting that genetically determined expression levels of mitochondrial structural regulators may influence kidney disease susceptibility and progression.

**Figure 1.**
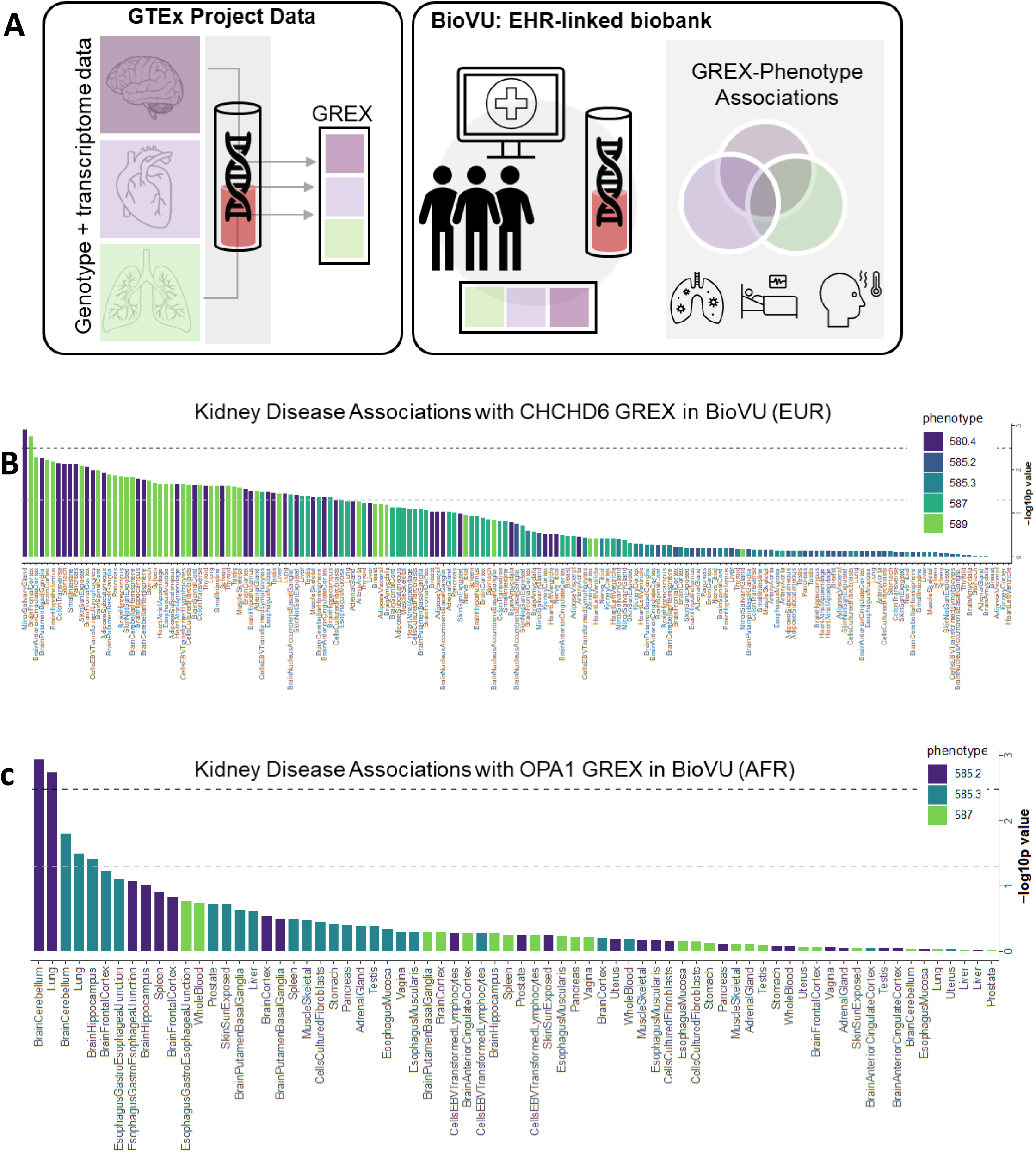
Assessing the relationship between the MICOS complex, genetically modeled gene expression, and kidney diseases in a clinical biobank. (A) Genetically-regulated gene expression (GREX) for *CHCHD3*, *CHCHD6*, and *OPA1* were calculated in BioVU participants (n=77,676) using models built from the GTEx version 8 data, which contain genotype data matched to transcriptome data from 838 donors across 49 tissues. (left) We identified cases and controls for kidney disease phenotypes from BioVU participants using ICD9/10 codes from Vanderbilt’s de-identified electronic health record database. (right) Genetically modeled expression was tested for association with kidney disease case status using logistic regression models, accounting for genetic ancestry (principal components/PC 1-10), sex, current age, median age of medical record, and genotype batch. (B) Association results for *CHCHD6* GREX and kidney phenotypes in individuals of European ancestry (EUR) are rank-ordered along the x-axis by decreasing -log10p value. Each bar represents a GREX model with tissues labeled on the x-axis. The bars are color-coded according to the kidney phenotype being tested. The black dotted line represents the within-tissue Bonferroni-corrected p-value, and the dotted grey line represents the nominal p-value (p=0.05). (C) Association results for OPA1 GREX and kidney phenotypes in individuals of African ancestry (AFR). Legend key: phecode 580.4: Renal sclerosis, NOS, phecode 585.2: Renal failure NOS, phecode 585.3: Chronic renal failure (CKD), phecode 587: Kidney replaced by transplant, phecode 589: Abnormal results of function study of kidney.

### Human Aging Causes Minimal Changes in Kidney Size

Previous studies have utilized magnetic resonance imaging (MRI) of solid renal masses as a proxy for pathologic classification and to define kidney structure^64,65^. Generally, it has been found that, after the age of 60, kidney volume decreases at approximately 16 cubic centimetres per decade ^66^. Thus, we utilized MRI to determine how the kidney is remodeled during the aging process. By enrolling female (Figures 2A & C) and male participants (Figures 2B & D), we created a “young” cohort (n = 14) consisting of individuals under 50 years old and an “old” cohort (n = 20) of individuals aged 60 or older. For both sexes, the kidney area did not show a significant change (Figures 2E and 2I). In-phase, which refers to aligned fat and water molecules, and out-of-phase, or opposed-phase, intensity were similarly minimally changed in both females and males across ageing (Figures 2F&G female, 2 J&K male). We observed that the old cohort of males had a significantly reduced in-phase intensity (Figure 2J). From there, we calculated the ratio of in-phase to out-of-phase intensity, which showed no significant difference (Figures 2H and L). When male and female subjects were combined, the kidneys from males and females were not significantly differentiated across the aging process. Interestingly, older females exhibited substantially lower cross-sectional areas than aged males, suggesting an increased susceptibility to aging with sex-dependent differences in kidney aging (Figure 1B).

**Figure 2.**
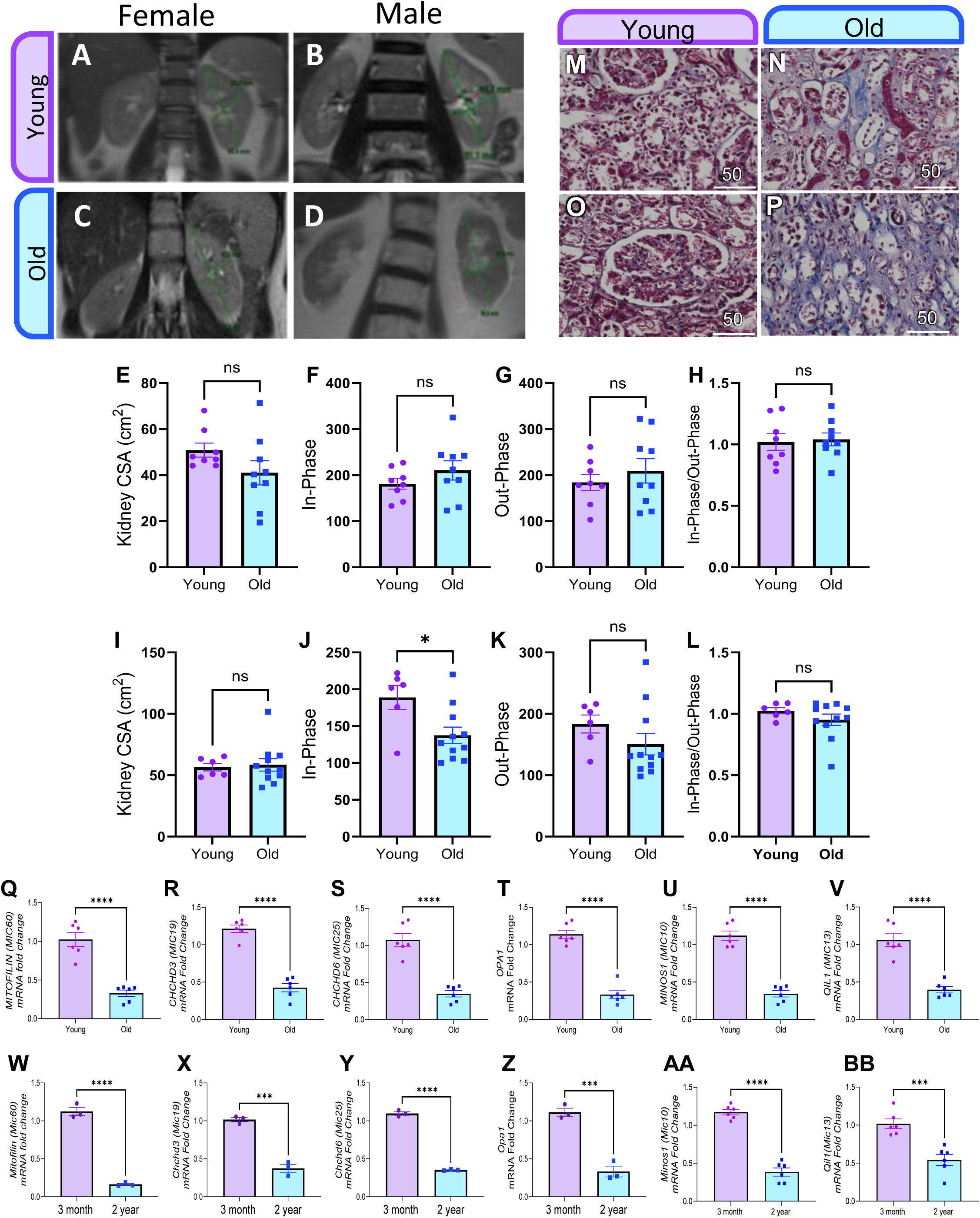
Age-associated structural, histological, and transcriptional alterations in the human and murine kidney. Cross-sectional MRI images of the left kidney from (A) females <50 years (14–42 years; n = 8), (B) males <50 years (14–48 years; n = 6), (C) females >60 years (65–90 years; n = 9), and (D) males >60 years (60–88 years; n = 11). Quantitative MRI analyses are shown separately for females (E–H) and males (I–L). (E, I) Left kidney cross-sectional area (CSA), calculated as the product of kidney length and width. (F, J) In-phase signal intensity, reflecting alignment of fat and water magnetic fields. (G, K) Out-of-phase signal intensity, reflecting misalignment of fat and water magnetic fields. (H, L) Ratio of in-phase to out-of-phase signal intensities. Representative histological images of human kidney cortex stained with Masson’s trichrome (M, O) and hematoxylin & eosin (N, P) from young adults (24–31 years; M, N) and older individuals (60–77 years; O, P). Scale bar, 50 µm. Relative mRNA expression of MICOS complex components in human kidney cortex from young (20–48 years) and old (65–90 years) individuals: (Q) *MIC60*/*MITOFILIN*, (R) *CHCHD3*/*MIC19*, (S) *CHCHD6*/*MIC25*, (T) *OPA1*, (U) *MINOS1*/*MIC10*, and (V) *QIL1*/*MIC13*. Relative mRNA expression of MICOS complex components in murine kidney cortex from 3-month-old and 2-year-old mice: (W) *Mic60*/*Mitofilin*, (X) *Chchd3*/*Mic19*, (Y) *Chchd6*/*Mic25*, (Z) *Opa1*, (AA) *Minos1*/*Mic10*, and (BB) *Qil1*/*Mic13*. Gene expression was assessed by quantitative PCR, normalized to housekeeping genes, and is presented as mean ± SEM. Statistical significance is indicated as ns, not significant; *p* < 0.05; \*\**p* < 0.001; \*\*\**p* < 0.0001.

Histological examination of human kidney cortex samples revealed distinct structural differences between age groups. Masson’s Trichrome staining demonstrated altered extracellular matrix composition and fibrotic changes in older individuals (60-77 years) compared to younger adults (24-31 years). (Figure 2M and N respectively). Complementary H&E staining further elucidated age-associated alterations in tissue architecture, cellular morphology, and organization within the kidney cortex (Figure 2O and P, young and old, respectively).

### Aging is Associated with Coordinated Downregulation of MICOS Components and Cristae Regulators in Human and Murine Kidneys

To determine whether aging is associated with altered expression of mitochondrial inner membrane organizing factors *in vivo*, we analyzed transcript levels of MICOS components and key cristae regulators in human kidney samples and in an independent murine aging model. In human kidneys, aging was associated with a robust and coordinated reduction in the expression of multiple MICOS subunits and cristae-associated genes. Quantitative mRNA analysis revealed a significant decrease in *MIC60(Mitofilin)* expression in aged samples compared with young controls (Figure 2Q). Similarly, transcripts encoding *CHCHD3(MIC19)* and *CHCHD6(MIC25)* were markedly reduced with age (Figure 2R, S). In addition to MICOS core components, expression of the mitochondrial inner membrane fusion regulator *OPA1* was significantly lower in aged human kidneys (Figure 2T). Transcripts corresponding to additional MICOS-associated subunits, including *MIC10(MINOS1)* and *MIC13(QIL1),* were also significantly downregulated in aged samples (Figure 2U, V), indicating a broad age-dependent suppression of the MICOS transcriptional program. Although we could not confirm that subjects had kidney disease, these results suggest a slight age-related decline in kidney structure. Although gross morphological changes may be minimal, we sought to elucidate other tissue changes that occur with aging.

To assess whether this age-associated MICOS decline is conserved across species, we next examined renal gene expression in a murine aging model comparing young (3-month-old) and aged (2-year-old) mice. Consistent with the human data, aged mouse kidneys exhibited a significant reduction in *Mitofilin (Mic60)* mRNA levels relative to young animals (Figure 2W). Expression of *Chchd3 (Mic19)* and *Chchd6 (Mic25)* was similarly decreased with age (Figure 2X, Y). In parallel, *Opa1* transcript levels were significantly reduced in aged mice (Figure 2Z), accompanied by decreased expression of *Minos1(Mic10)* and *Quil1(Mic13)* (Figure 2AA, BB).

### Murine Aging Results in Interstitial Fibrosis and Oxidative Stress

Previous studies have shown that interstitial fibrosis on kidney biopsy is a prognostic indicator and can represent nephropathy, although its diagnostic effectiveness is mixed^67,68^. We examined young (4–5 months old; Figure 3A) and old (21–23 months old; Figure 3B) C57BL/6J mice using trichrome blue to stain connective tissue. Concurrent with previous studies^55^, we found that the trichrome area percentage was higher in the old mice (11.0%) than in the young mice (5.8%), indicating that older mice had a greater degree of interstitial fibrosis (Figure 3C). From there, we used immunohistochemistry to examine nitrotyrosine, a biomarker of oxidative stress^69^, as shown in the brown areas (Figure 3A’ & 3B’). Studies have shown that increased nitrotyrosine levels correlate with renal dysfunction and inflammatory processes, serving as a biomarker for kidney diseases such as AKI and CKD, as well as overall mortality in these disease states^70–72^. Immunohistochemistry revealed a significant increase in nitrotyrosine in tubular epithelial cells and podocytes of old mice compared with their young counterparts (Figure 3D). This indicates that oxidative stress occurs with aging, so we sought to understand how mitochondrial structure also changes.

**Figure 3.**
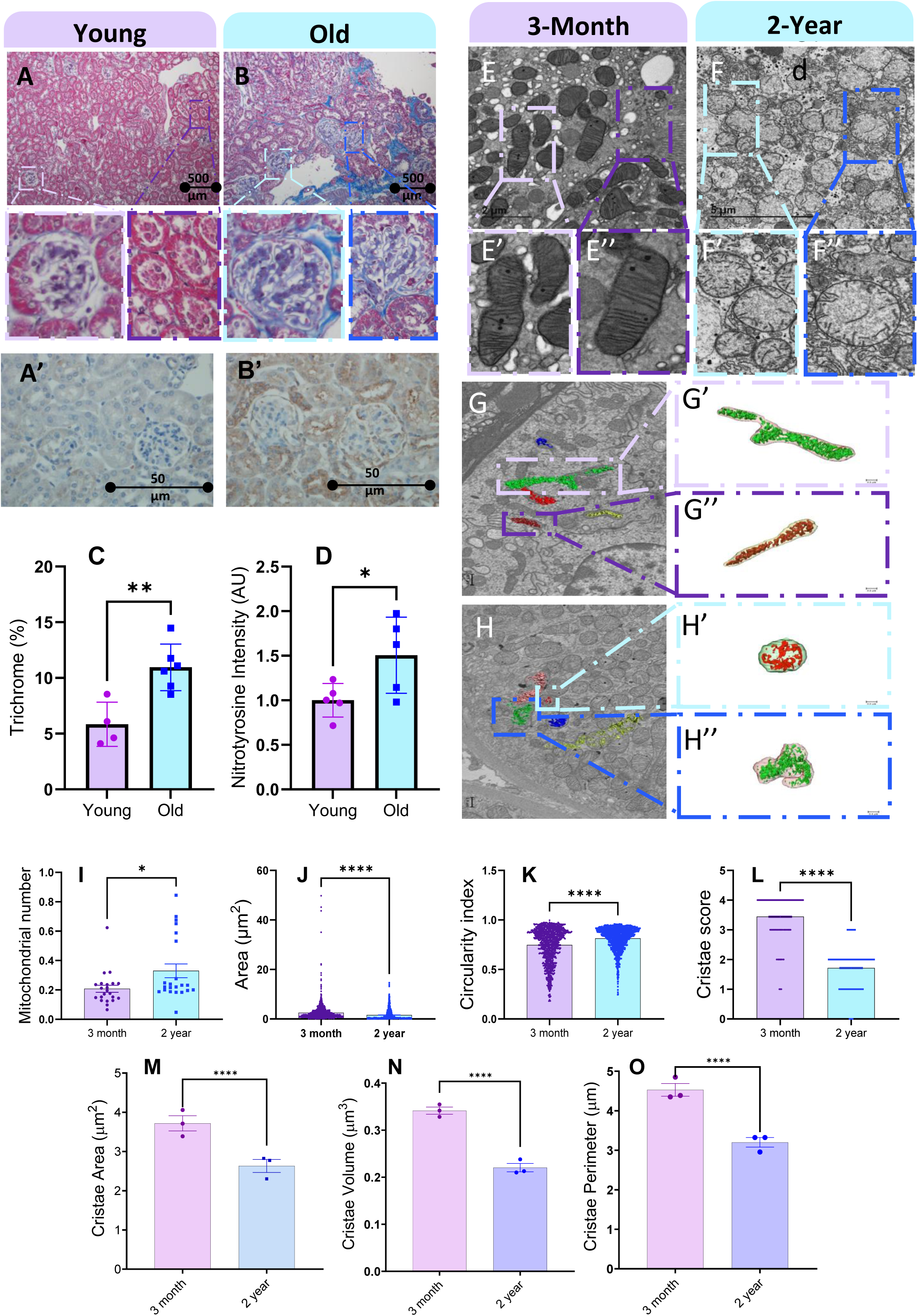
Age-associated histological and ultrastructural remodeling of renal mitochondria in mice. (A, B) Representative images of Masson’s trichrome staining of the kidney cortex from young (4–5 months) and old (21–23 months) mice. Insets show higher-magnification views of glomerular and tubular regions. (A′, B′) Representative immunohistochemistry for nitrotyrosine (brown) in the kidney cortex from young (4–5 months) and old (21–23 months) mice. (C) Quantification of trichrome-positive area (%) and (D) nitrotyrosine staining intensity corresponding to panels A′ and B′. (E–E″) Representative transmission electron microscopy (TEM) images of kidney tissue from 3-month-old male mice and (F–F″) 2-year-old male mice, showing mitochondrial ultrastructure at increasing magnifications. (G, H) Representative serial block-face scanning electron microscopy (SBF-SEM) images illustrating three-dimensional cristae morphology in 3-month-old and 2-year-old samples, respectively. Boxed regions indicate mitochondria selected for cristae segmentation and magnified in (G′–G″) and (H′–H″). Quantifications in panels I–L were derived from TEM images, whereas cristae morphometric parameters in panels M–O were obtained from three-dimensional SBF-SEM reconstructions. (I) Quantification of mitochondrial number (J), mitochondrial area (K), circularity index (L), and cristae score, a semiquantitative measure of observed cristae organization. (M) Cristae area, (N) cristae volume, and (O) cristae perimeter derived from three-dimensional reconstructions. Mitochondrial measurements were obtained from approximately 1,050 mitochondria at 3 months and approximately 1,450 mitochondria at 2 years. Cristae score analysis included 1,093 mitochondria. Each dot represents an individual mitochondrion, with variable numbers per animal. For TEM-based analyses, each age group comprised 3 mice. Data are presented as mean ± SEM. Statistical comparisons were performed using Mann–Whitney tests. Statistical significance is denoted as ns (not significant), p < 0.05, p < 0.01, and **p ≤ 0.0001.

### Ultrastructural Analysis Reveals Age-Associated Enlargement and Rounding of Renal Mitochondria

Previous studies have shown that aging influences mitochondrial dynamics and morphology in the kidney^73^. To assess age-dependent ultrastructural changes, we analyzed TEM images of renal tubular mitochondria from 3-month- and 2-year-old mice (Figure 3E–F″), using samples obtained from three animals per group. Examination of TEM sections revealed clear alterations in mitochondrial morphology with advancing age. Quantitative morphometric analysis demonstrated a significant increase in average mitochondrial area in aged kidneys. Mitochondria from 3-month-old mice exhibited a mean area of 1.57 µm² ± 2.09 µm² (SD), whereas mitochondria from 2-year-old mice showed a larger mean area of 2.54 µm² ± 3.33 µm² (SD) (Figure 3J). In parallel, mitochondrial shape was altered with aging, as reflected by a significant increase in the circularity index from 0.746 ± 0.174 (SD) in 3-month-old mice to 0.813 ± 0.116 (SD) in 2-year-old mice (Figure 3K). In contrast, mitochondrial number did not differ significantly between young and aged kidneys (Figure 3I).

Together, these findings indicate that aging is associated with mitochondrial enlargement and increased rounding in renal tubular cells. While increased mitochondrial size could theoretically expand membrane surface area, such morphological changes are also consistent with pathological mitochondrial swelling, a feature commonly linked to mitochondrial stress and dysfunction^74^. Given that mitochondrial ultrastructure, including cristae organization, plays a critical role in regulating bioenergetics^75^, these age-associated morphological alterations prompted further examination of cristae integrity.

### Age-Dependent Loss of Cristae Integrity Revealed by Cristae Scoring and Quantitative Morphometric Analysis

Ultrastructural analysis of TEM images revealed marked age-associated disruption of cristae organization in renal tubular mitochondria (Fig. 3E″, F″, magnified regions). To quantitatively assess these changes, we applied a previously established cristae scoring system^62, 76, 77^, an ordinal scale ranging from 0 to 4 that reflects progressive degrees of cristae organization, from little-to-no discernible cristae (score 0) to well-formed, densely packed cristae (score 4). Cristae scoring revealed a significant decline in cristae integrity with aging. Mitochondria from 3-month-old mice predominantly exhibited well-organized cristae, with a mean cristae score of 3.44 ± 0.794 (SD), whereas mitochondria from 2-year-old mice showed substantially fewer well-formed cristae, with a reduced mean score of 1.71 ± 0.693 (SD) (Fig. 3L). These results indicate a pronounced loss of cristae structural integrity in aged kidneys.

To complement this qualitative scoring and obtain objective measurements of cristae architecture, we performed morphometric analyses of individual cristae from TEM images. Quantitative assessment revealed a significant reduction in cristae area in aged mitochondria compared with young samples (Fig. 3M). In parallel, cristae volume was markedly decreased in 2-year-old mice relative to 3-month-old controls (Fig. 3N), indicating a substantial loss of inner membrane surface area at the level of individual cristae. Consistent with these findings, cristae perimeter was also significantly reduced with aging (Fig. 3O), reflecting diminished cristae length and reduced architectural complexity.

Together, cristae scoring, quantitative TEM-based morphometrics, and complementary SBF-SEM analyses consistently demonstrate that aging is associated with a profound loss of cristae integrity, size, and complexity in renal mitochondria. Importantly, these defects occur despite an overall increase in mitochondrial size, indicating a decoupling between mitochondrial enlargement and preservation of inner membrane architecture during renal aging.

### SBF-SEM Reveals Aging Results in Reduced Mitochondrial Volume in Kidney Tissue

With these observations, we aimed to determine whether mitochondrial networks undergo changes in response to aging. In Figure 4, we show representative orthoslice images of the kidney tissue at each aging point (Figures 4A, B), the overlay of the 3-D reconstruction on orthoslice (Figures 4A’-B’), and the isolated 3-D reconstruction (Figures 4A“-B”), with each color representing an independent mitochondrion. We found that the average mitochondrial 3D area, a measure of outer mitochondrial membrane surface area, did not significantly change between 3-month (mean 8.29 µm^2^ ± 10.1 µm^2^ SD) versus 2-year (6.46 µm ± 5.31 µm SD) cohorts, unlike our TEM findings, despite great interindividual heterogeneity (Figures 4C and F). However, the perimeter decreased between the 3-month (mean 14,328 µm ± 17,723 µm) and 2-year (10,241 µm ± 8,273 µm SD) cohorts, and the 2-year cohort also showed less intra-individual heterogeneity (Figures 4D and G). This trend towards smaller mitochondria was reflected in the comparison of volume between the 3-month (0.920 µm^3^ ± 1.06 µm^3^ SD) and 2-year (0.741µm^3^ ± 0.695 µm^3^ SD) cohorts (Figures 4E and J). Quantitative 3D analysis revealed no significant age-associated change in mitochondrial sphericity between 3-month- and 2-year-old mice (Figure 4H). Mitochondrial complexity index was significantly reduced in 2-year-old mice compared with 3-month-old controls, indicating a loss of structural intricacy with aging (Figure 4I).

**Figure 4.**
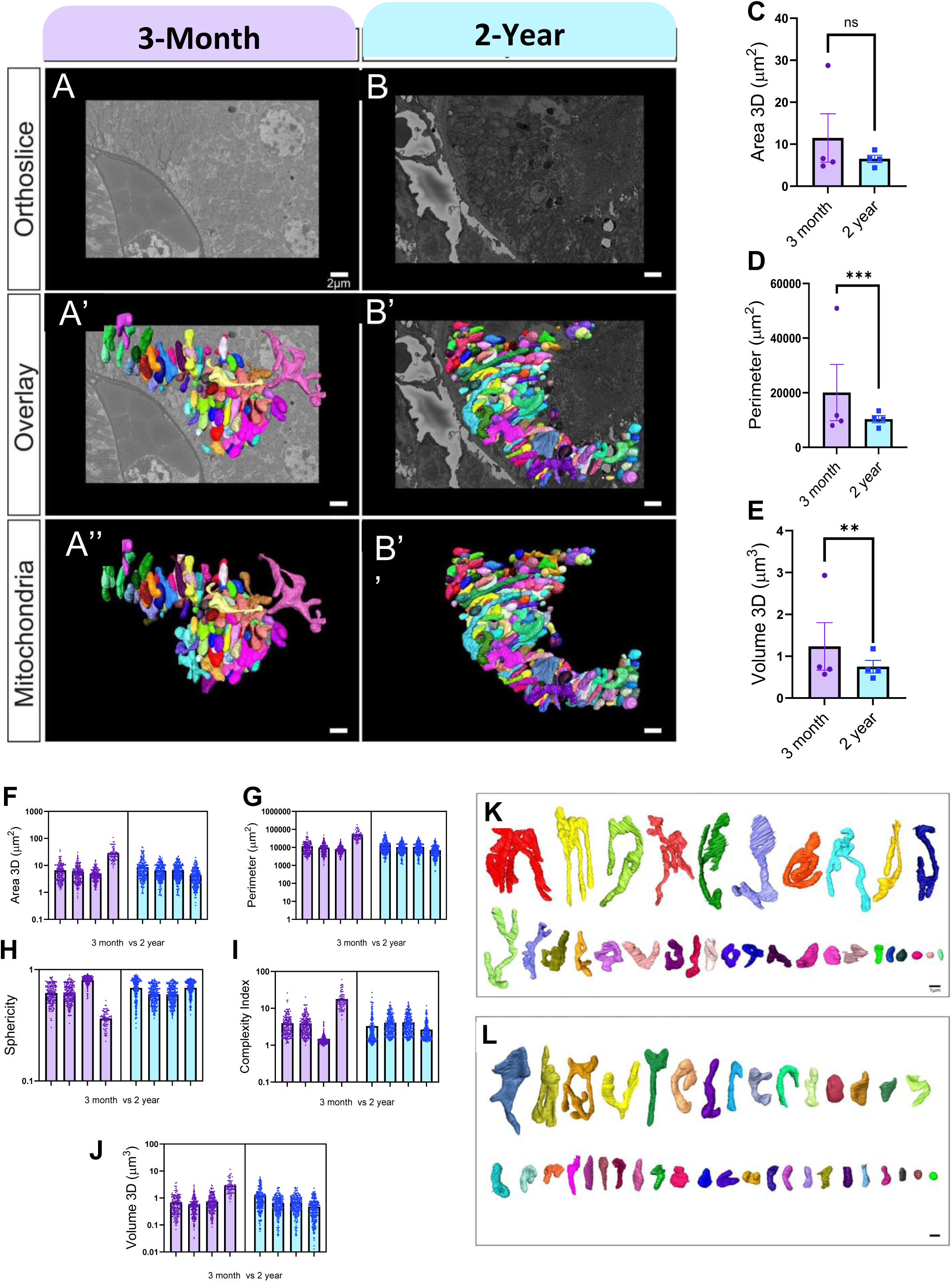
Three-dimensional mitochondrial morphometry in the aging mouse kidney assessed by SBF-SEM. (A, B) Representative SBF-SEM orthogonal (ortho) slices of kidney tissue from 3-month-old and 2-year-old male mice. (A′, B′) Three-dimensional reconstructions of mitochondria from 3-month-old and 2-year-old kidney tissue, respectively, overlaid on corresponding ortho slices. (A″, B″) Isolated three-dimensional reconstructions of individual mitochondria from 3-month-old and 2-year-old samples for enhanced visualization. Quantitative analyses of three-dimensional mitochondrial reconstructions are shown as follows: (C, F) mitochondrial 3D area, (D, G) mitochondrial perimeter, (E, J) mitochondrial volume, (H) sphericity, and (I) complexity index. Each dot represents the average value from a single mouse, with a variable number of mitochondria analyzed per mouse (n = 4 mice per age group; 83–251 mitochondria per mouse). In total, 740 mitochondria from 3-month-old mice and 962 mitochondria from 2-year-old mice were included for statistical analysis. Data are presented as mean ± SEM. Statistical comparisons were performed using the Mann–Whitney test, with significance denoted as ns (not significant), **p ≤ 0.01, and ***p ≤ 0.001. (K, L) Mito-typing of reconstructed mitochondria from 3-month-old (K) and 2-year-old (L) mice, arranged by mitochondrial volume to highlight qualitative differences in mitochondrial morphology.

To further elucidate age-related changes and characterize the mitochondrial types in each age cohort, we employed mito-otyping, a method similar to karyotyping, to organize mitochondria based on their volumes, thereby enhancing the visualization of overall mitochondrial diversity (Figure 4K and L). Critically, this approach revealed few significant morphological changes, with only branching showing reductions. In combination, the aged kidney mitochondrial morphology resembled that of healthy mitochondria, with a reduced size and lacking a phenotype or fragmentation. Since mitochondria showed a variety of structural changes due to aging, we turned our attention to the MICOS complex as a potential mechanistic regulator of these age-related changes.

### Three-Dimensional SBF-SEM Analysis Reveals Age-Associated Reduction in Mitochondrial Complexity and Mitochondria–ER Contact in Mouse Kidney

To examine age-related changes in mitochondrial architecture in kidney proximal tubule cells, we performed high-resolution three-dimensional imaging using serial block-face scanning electron microscopy (SBF-SEM) on tissue from young (3-month-old) and aged (2-year-old) mice (Fig. 5A–D). Volumetric datasets spanning 50 µm × 10 µm × 10 µm were acquired, and individual mitochondria were segmented across 25 consecutive serial sections to generate three-dimensional reconstructions (Fig. 5C, D). Also, due to the substantial time demands of 3D reconstruction and males’ higher kidney disease burden from sex differences^79–81^, our study used only a male model.

**Figure 5.**
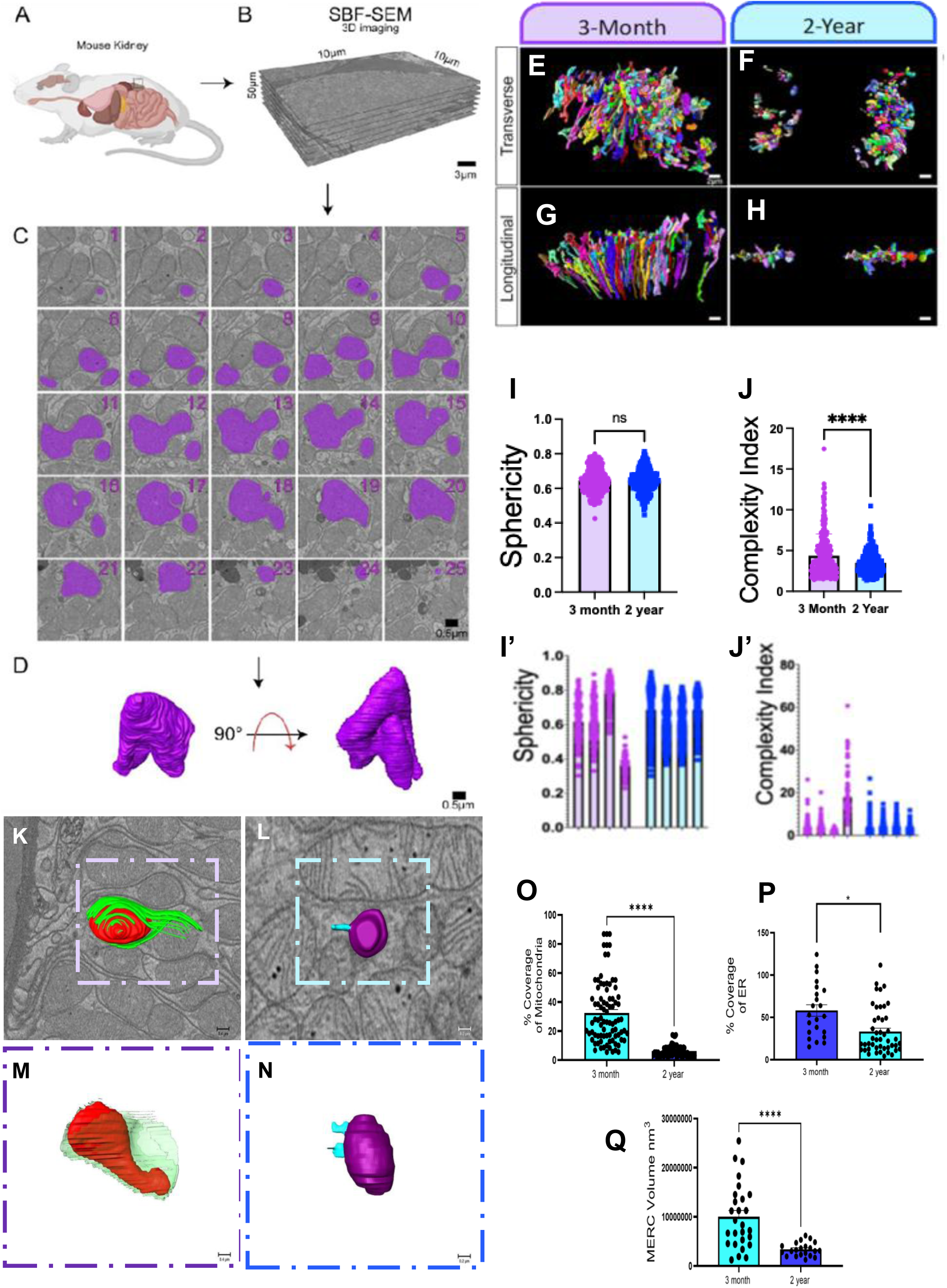
Three-dimensional analysis of mitochondrial morphology and mitochondria–ER contacts in the aging mouse kidney using SBF-SEM. (A) Schematic of the experimental workflow illustrating kidney tissue collection from young (3-month-old) and aged (2-year-old) mice. (B) SBF-SEM was used to acquire volumetric datasets (50 µm × 10 µm × 10 µm) for three-dimensional reconstruction. (C) Representative two-dimensional SBF-SEM serial sections (25 consecutive slices) from proximal tubule cells, highlighting a segmented mitochondrion (purple). Scale bar, 0.5 µm. (D) Three-dimensional reconstruction of the mitochondrion shown in (C) with a 90° rotation, illustrating complex membrane topology. Scale bar, 0.5 µm. (E, F) Three-dimensional reconstructions of mitochondria from 3-month-old (E) and 2-year-old (F) mouse kidneys are shown in transverse orientation. (G, H) Corresponding longitudinal views of mitochondrial reconstructions from 3-month-old (G) and 2-year-old (H) samples. Each color represents an individual mitochondrion. Scale bars, 0.5 µm. (I) Quantification of mitochondrial sphericity shows no significant difference between young and aged groups. (J) Quantification of the mitochondrial complexity index demonstrates a significant reduction in aged kidneys compared with young controls. (I′, J′) Distribution of sphericity and complexity index values for individual mitochondria, illustrating greater morphological heterogeneity in young samples and an overall reduction in complexity with aging. (K, L) Representative overlays of three-dimensional reconstructions of mitochondria and wrapping endoplasmic reticulum (wrappER) on orthogonal SBF-SEM slices from (K) 3-month-old and (L) 2-year-old kidneys. (M, N) Side views of three-dimensional reconstructions of mitochondria and associated wrappER from (M) 3-month-old and (N) 2-year-old samples. (O) Quantification of the percentage coverage of mitochondria by ER in 3-month-old and 2-year-old mice. (P) Quantification of ER coverage in young and aged kidneys. (Q) Quantification of mitochondria–ER contact (MERC) volume in 3-month-old and 2-year-old mice. Data are presented as mean ± SEM. Individual data points represent independent mitochondria reconstructed from SBF-SEM volumes. Statistical comparisons between age groups were performed using a two-tailed Mann–Whitney U test. Statistical significance is denoted as ns (not significant), p < 0.05, and ***p < 0.0001.

Three-dimensional reconstructions revealed clear differences in mitochondrial organization between young and aged kidneys. In 3-month-old mice, mitochondria exhibited elongated and highly branched morphologies in both transverse and longitudinal orientations (Fig. 5E, G). In contrast, mitochondria from 2-year-old mice appeared more fragmented and less elaborately branched (Fig. 5F, H). Quantitative analysis of mitochondrial sphericity demonstrated no significant difference between young and aged groups (Fig. 5I, I′), indicating that overall mitochondrial roundness is preserved with aging. In contrast, quantification of the mitochondrial complexity index revealed a significant reduction in aged kidneys compared with young controls (Fig. 5J, J′), consistent with a loss of three-dimensional structural complexity. Distribution analysis further showed greater heterogeneity and higher complexity values in mitochondria from young kidneys, whereas mitochondria from aged kidneys clustered toward lower complexity values.

To assess age-associated changes in mitochondria–endoplasmic reticulum (ER) interactions, we examined three-dimensional overlays of mitochondria and wrapping ER (wrappER) on orthogonal SBF-SEM slices. In young kidneys, mitochondria frequently displayed extensive ER coverage consistent with a wrappER phenotype (Fig. 5K, M). In contrast, mitochondria from aged kidneys showed a visibly reduced association with the ER (Fig. 5L, N). Quantitative analysis confirmed a significant reduction in the percentage of mitochondrial surface covered by ER in aged mice compared with young controls (Fig. 5O). Similarly, overall ER coverage within the analyzed volumes was reduced in aged kidneys (Fig. 5P). Consistent with these findings, quantification of mitochondria-ER contact (MERC) volume revealed a significant decrease in aged mice relative to young animals (Fig. 5Q). The wrappER-associated morphology observed in young kidneys resembles ER-mitochondria interaction phenotypes previously described in other tissues^78^.

### Global Metabolic and Lipid Profiling Highlights Dynamic Changes in the Aged Kidney

Following our observations of age-associated mitochondrial structural dysregulation, we sought to define how these morphological changes impact metabolic homeostasis in the kidney. To this end, we performed comprehensive metabolic and lipid profiling of young and aged kidney tissues. Metabolic profiling revealed broad disruptions in amino acid metabolism, nucleotide biosynthesis, and redox cofactors in aged kidneys (Figures 6A-R). Amino acids are central to mitochondrial function, supporting energy production, gluconeogenesis, nitrogen metabolism, antioxidant defense, and specialized mitochondrial pathways^83^. In aged kidneys, we detected significant reductions in methionine, valine, threonine, leucine, isoleucine, and glycine. Notably, glycine, serine, and threonine metabolism—key components of one-carbon metabolism—were markedly perturbed (Figures 6J, 6M). These pathways are tightly linked to mitochondrial function and purine biosynthesis, and disruptions may reflect impaired flux through the mitochondrial glycine cleavage system (GCS), influencing synthesis of purines, pyrimidines, and other small molecules (Figures 6D-R). Consistent with this, we observed significant declines in purine intermediates, including inosine monophosphate (IMP), uridine monophosphate (UMP), ribose-phosphate, and related purine metabolites, suggesting compromised nucleotide biosynthetic capacity in aged kidneys (Figure 6D-G). Surprisingly, however, total adenylate pools (AMP, ADP, and ATP) were preserved with age (Figure 6A-C). These findings indicate that despite upstream perturbations in nucleotide and amino acid metabolism, steady-state ATP levels remain buffered.

**Figure 6.**
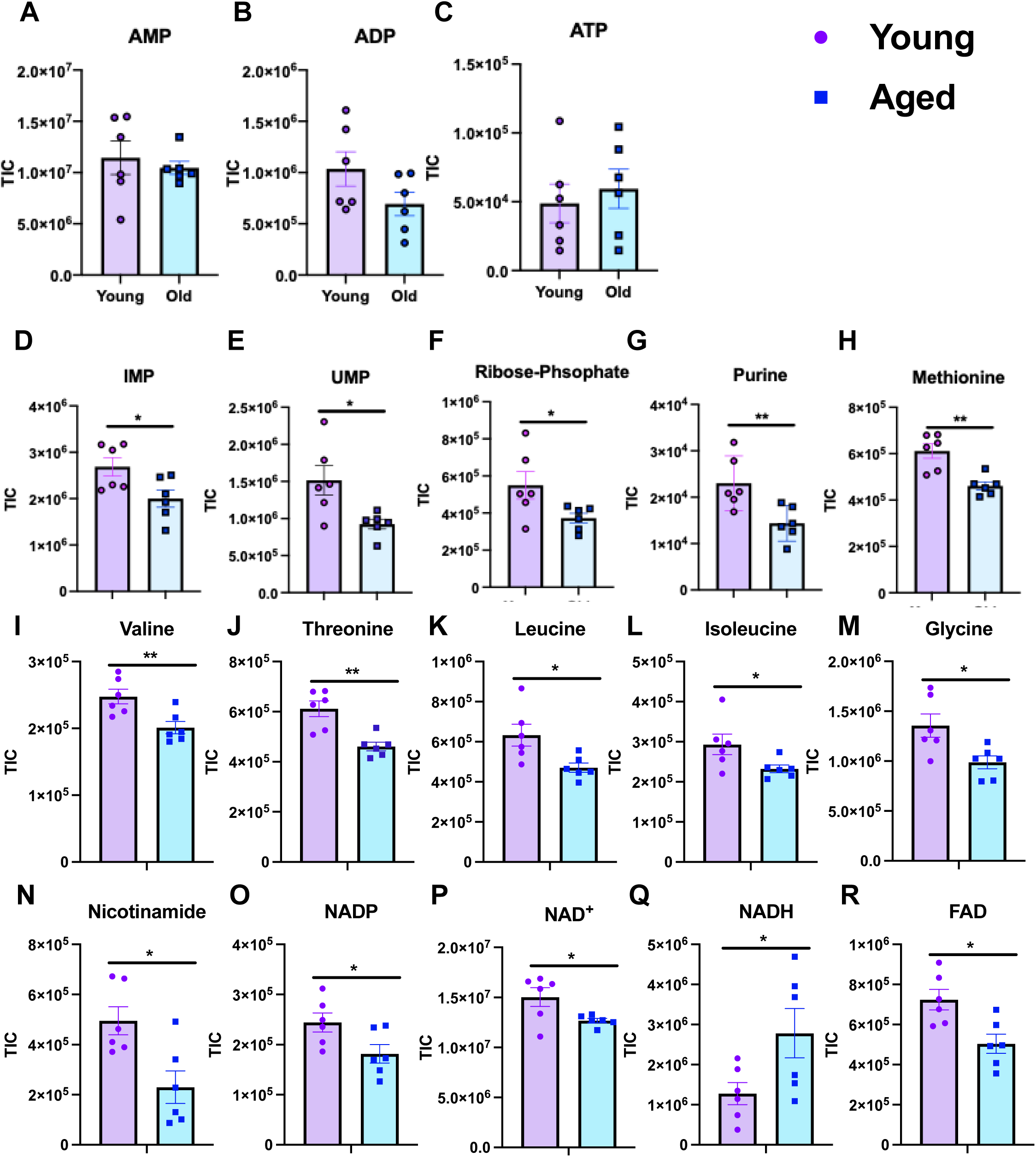
Targeted LC–MS profiling reveals age-associated metabolic remodeling in kidney tissues. (A–R) Targeted metabolite pool sizes highlighting pathways altered with age, including purine metabolism, amino acid metabolism/biosynthesis, and redox/NAD⁺ metabolism. (A–C) Adenylate pools (AMP, ADP, ATP) show no significant age-related changes. (D–G) Purine metabolism intermediates, including IMP, UMP, ribose-phosphate, and purine, are significantly reduced in aged kidneys. (H–M) Amino acid metabolites, including methionine, valine, threonine, leucine, isoleucine, and glycine, are decreased with age. (N–R) Redox and NAD⁺ metabolism intermediates, including nicotinamide, NADP, NAD⁺, NADH, and FAD, demonstrate age-associated imbalance characterized by reduced NAD⁺, NADP, and FAD levels and increased NADH. Metabolite abundances are presented as total ion counts (TIC), as indicated on the y-axes. Young, n = 6; aged, n = 6. Data are shown as mean ± SEM. Statistical significance was determined using Student’s t-test; *p < 0.05, **p < 0.01.

To further interrogate mitochondrial energy metabolism, we examined redox cofactors. Aged kidneys exhibited significant declines in NAD⁺, NADP, nicotinamide (NAM), and FAD levels, alongside an increase in NADH. This shift suggests an imbalance in redox state consistent with reductive stress. Importantly, although total adenylate pools were unchanged, the altered NAD⁺/NADH ratio and decreased FAD indicate impaired oxidative capacity and electron carrier availability. Together, these findings suggest two non-mutually exclusive interpretations: (1) aged kidneys engage compensatory mechanisms that preserve ATP levels despite upstream metabolic remodeling, or (2) metabolic rewiring follows a temporal hierarchy, wherein redox imbalance and nucleotide biosynthetic deficits precede overt ATP depletion. These metabolic signatures align with the mitochondrial morphological and transcriptional alterations observed in our imaging and gene expression analyses, supporting a model in which structural mitochondrial remodeling drives redox and biosynthetic perturbations prior to energetic collapse. Importantly, these measurements were performed in bulk kidney tissue. Thus, compartment-specific alterations—particularly within mitochondrial ATP pools—may be masked. We cannot definitively conclude whether mitochondrial-localized ATP production is preserved, as cytosolic buffering mechanisms may obscure organelle-specific deficits.

We profiled the kidney lipidome in young and old mice to assess age-associated differences in lipid composition. Principal component analysis showed modest structure by age, with a partial shift in the distribution of samples along the first two components but substantial overlap between groups. Consistent with this, differential abundance analysis at the level of individual lipid species did not reveal a clear statistical signal after correction for multiple testing, with adjusted p-values remaining above conventional significance thresholds across the dataset (Supplementary Table 1, Figure Volcano), indicating that age-related differences are not driven by large changes in a small number of individual molecules.

At the level of lipid classes and structural groupings, lipid set enrichment analysis did identify some coordinated differences between the groups (Enrichment Figures, Supplementary Table 2). Cardiolipins (CL) were positively enriched in young kidneys (ES = 0.465, padj = 3.22×10⁻⁴), along with related mitochondrial lipid sets including dilysocardiolipins (DLCL; ES = 0.737, padj = 0.0071) and lysophosphatidylinositols (LPI; ES = 0.909, padj = 0.0084). Triglycerides (TG; ES = −0.360, padj = 1.91×10⁻⁵) and N-acyl ethanolamides (NAE; ES = −0.801, padj = 2.05×10⁻⁵) were negatively enriched, indicating relatively higher representation in older kidneys (Supplementary Table). With respect to structural features, several chain length–defined bins were significantly enriched in young kidneys, including total_cs_8 (ES = 0.531, padj = 3.22×10⁻⁴), total_cl_60 (ES = 0.708, padj = 0.0046), and total_cl_62 (ES = 0.757, padj = 0.0046), while shorter-chain groupings showed relative depletion.

Lipidomic profiling of young and aged kidney tissues further revealed age-dependent remodeling of lipid classes and acyl chain composition (Figures 7A–C). In aged kidneys, significant alterations were observed in triglyceride oligomers (TGO), triglycerides (TG), sterols (ST), N-acylethanolamines (NAE), lyso-phosphatidyl-inositol (LPI), dihexosylceramides (Hex2Cer), dilysocardiolipin (DLCL), and cardiolipin (CL) compared to young controls (Figure 7B). Notably, cardiolipin and dilysocardiolipin are critical structural phospholipids of the inner mitochondrial membrane that stabilize respiratory chain supercomplexes and maintain cristae architecture. Age-associated alterations in cardiolipin abundance and acyl chain composition likely impair electron transport chain (ETC) organization and efficiency. Because FAD and NAD⁺ function as essential electron carriers feeding into the ETC, structural membrane perturbations may reduce effective electron flux, contributing to the observed decline in NAD⁺ and FAD pools and accumulation of NADH. Thus, mitochondrial membrane remodeling in aged kidneys provides a structural framework that may underlie the redox imbalance consistent with reductive stress. In addition, significant differences in lipid chain length with age (Figure 7C) suggest altered membrane fluidity and curvature, which can further disrupt mitochondrial bioenergetics, calcium handling, and reactive oxygen species signaling. Together, these lipidomic changes support a model in which age-dependent membrane remodeling compromises mitochondrial respiratory efficiency, reinforcing the redox and metabolic rewiring observed in our metabolic profiling analysis.

**Figure 7.**
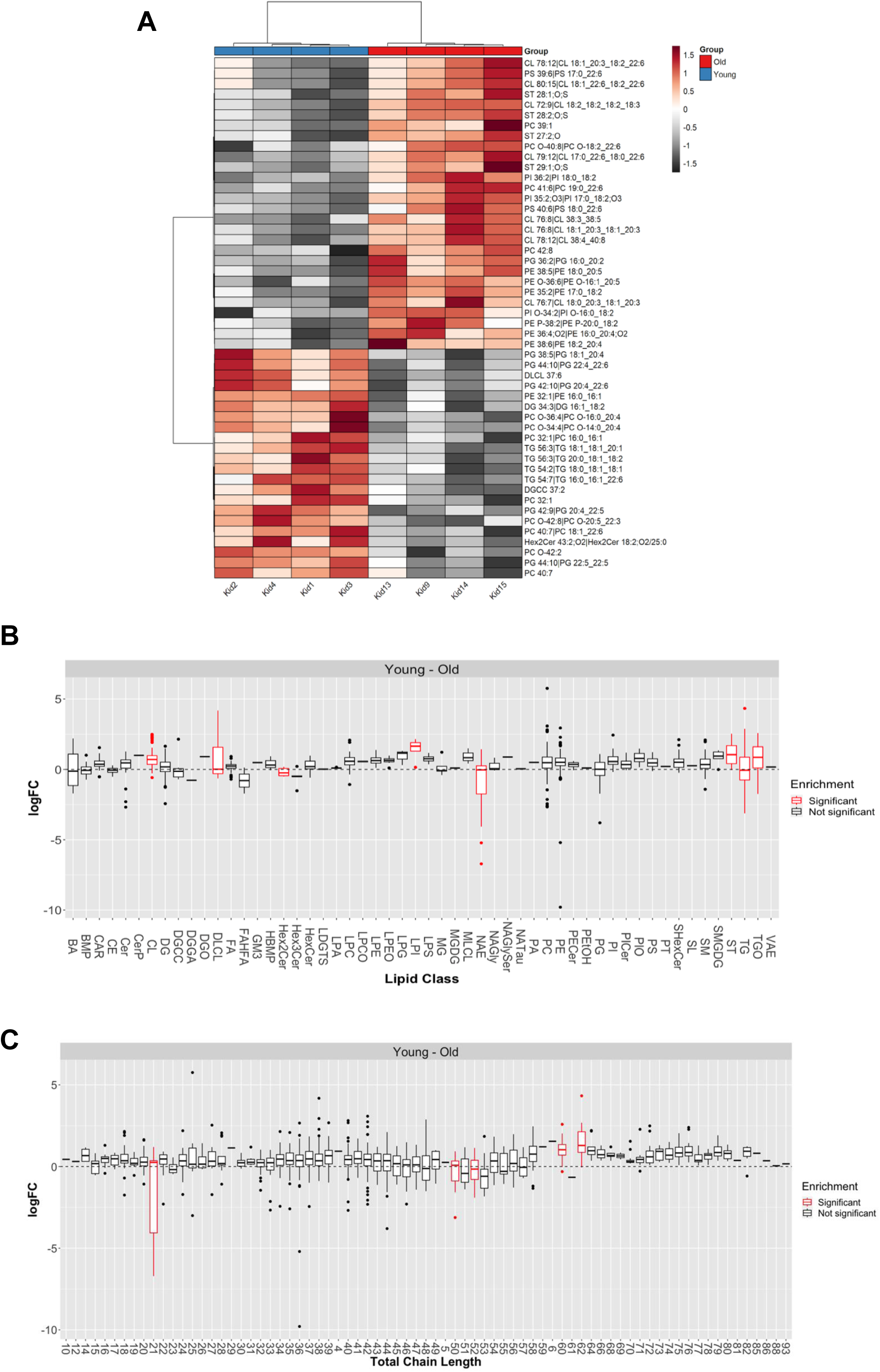
Aging drives coordinated remodeling of lipid classes and acyl chain composition in the mouse kidney. (A) Heatmap depicting the relative abundance of lipid species in kidney tissue from young and aged mice. Lipid intensities were normalized across samples and log₂-transformed prior to hierarchical clustering. Columns represent individual biological replicates (young, *n* = 6; aged, *n* = 6), and rows represent individual lipid species grouped by class. The color scale indicates relative abundance (log₂ fold change), with red denoting higher abundance and grey to black denoting lower abundance relative to the dataset mean, as shown in the accompanying scale bar. The top annotation indicates the age group. (B) Lipid class enrichment analysis showing the distribution of log fold change (logFC; young minus old) for individual lipid species within each lipid class. Boxes represent the interquartile range with the median indicated; whiskers denote 1.5× interquartile range; individual points represent lipid species. Lipid classes highlighted in red indicate statistically significant enrichment after multiple-testing correction. (C) Lipid chain-length enrichment analysis showing logFC (young minus old) for lipid species grouped by total acyl chain length. Boxes and whiskers are defined as in (B). Chain lengths highlighted in red indicate statistically significant enrichment following multiple-testing correction. Significantly altered lipid classes and chain lengths were defined as those with false discovery rate (FDR)–adjusted *p* < 0.05 and absolute logFC > 1. Young and aged groups each consisted of *n* = 6 biological replicates. Data are presented as mean ± SEM where applicable. For metabolite-level comparisons, statistical significance was assessed using an unpaired two-tailed Student’s *t*-test with FDR correction for multiple comparisons. Statistical significance is denoted as p ≤ 0.05 (*) and p ≤ 0.01 (**).

### Loss of OPA1 or MICOS Components Disrupts Mitochondrial Ultrastructure and Morphology

Oxidative stress is a defining feature of kidney ageing and CKD, with mitochondria representing a major cellular source of reactive oxygen species through the electron transport chain^34,93,94^. Cristae architecture is a key determinant of mitochondrial respiratory efficiency and redox balance and is maintained by the mitochondrial contact site and cristae organizing system (MICOS)^86–89^. We previously observed an age-associated decline in transcripts encoding core MICOS components—*Mic60* (Mitofilin), *Chchd3* (MIC19), and *Chchd6* (MIC25)—and the mitochondrial fusion regulator *Opa1* in kidney tissue^84,85,41,90^. These observations prompted us to test whether loss of MICOS integrity, independently or in parallel with OPA1, is sufficient to disrupt mitochondrial ultrastructure and promote oxidative stress^91^.

Since the loss of *Opa1* triggers morphological changes, we used it as a positive control for morphological changes^90,92^. We performed siRNA deletions of *Chchd3*, *Mitofilin*, *Chchd6*, and *Opa1* in immortalized human embryonic kidney cells (HEK293 cells). From there, we performed TEM in each condition and compared them with a control (Figure 8A-E). As expected, *Opa1* deletion resulted in significant decreases in mitochondrial area, perimeter, and length, accompanied by an inverse increase in circularity index, which was expected due to impaired fusion dynamics (Figures 8F-I). *Chchd3* deletion exhibits a more drastic phenotype, characterized by a reduced mitochondrial area, whereas *Chchd6* deletion shows a smaller decrease compared to the control. *Mitofilin* deletion demonstrated no significant differences (Figure 8F). Interestingly, *Chchd3* deletion in HEK293 cells results in increased perimeter and length, despite decreased area and circularity index (Figures 8F-I). *Chchd3* KO cells had nearly completely elongated mitochondria, unlike those in *Opa1* deletion. Taken together, this suggests that, while the MICOS complex KO phenotype is distinct from OPA1 loss, it can, beyond its canonical roles in cristae integrity, also modulate mitochondrial structure^85^. Since cristae and mitochondrial dysfunction were hallmarks of aging kidneys, we sought to understand the further functional implications of the age-dependent loss of the MICOS complex.

**Figure 8.**
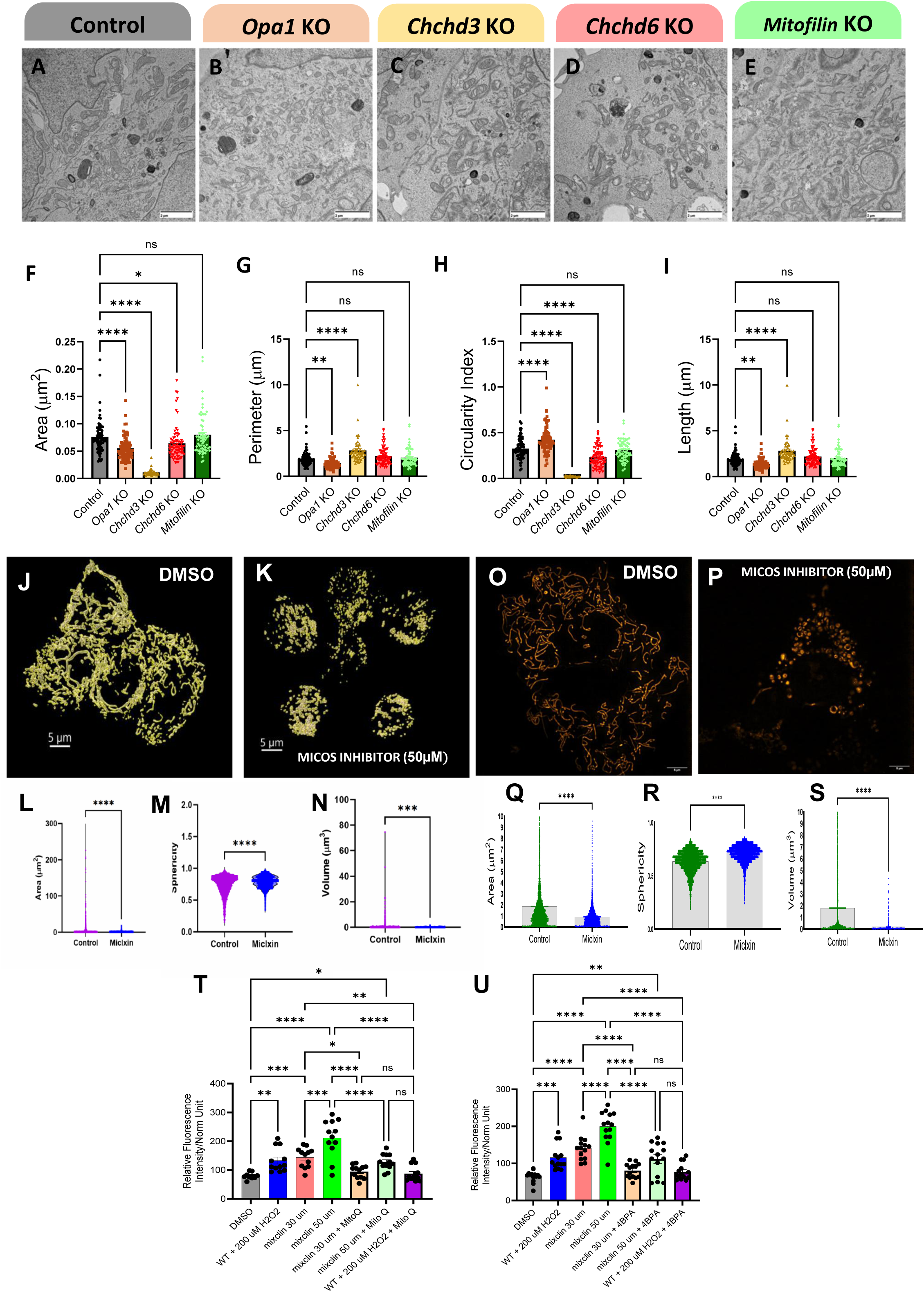
Genetic and pharmacological disruption of MICOS–OPA1 signaling remodels mitochondrial architecture across models and alters redox homeostasis *in vitro*. (A–E) Representative transmission electron microscopy (TEM) images of mitochondria from kidney tissue of control, *Opa1* knockout (KO), *Chchd3* KO, *Chchd6* KO, and *Mitofilin* KO samples. Control mitochondria display intact cristae organization, whereas loss of OPA1 or MICOS components results in disrupted cristae architecture and altered mitochondrial morphology. Scale bars as indicated. (F–I) Quantitative morphometric analysis of mitochondrial ultrastructure corresponding to panels A–E, including mitochondrial area (F), perimeter (G), circularity index (H), and length (I). Each data point represents an individual mitochondrion; bars indicate mean ± SEM. Statistical significance is denoted as ns (not significant), p ≤ 0.05, p ≤ 0.01, and **p ≤ 0.0001. (J–S) Pharmacological perturbation of MICOS in cultured cells recapitulates mitochondrial fragmentation phenotypes observed in kidney tissue. Representative three-dimensional mitochondrial reconstructions from HEK293 cells (J, K) and COS7 cells (O, P) treated with vehicle control (DMSO) or the MICOS inhibitor Miclixin (50 µM). Vehicle-treated cells exhibit interconnected mitochondrial networks, whereas MICOS inhibition induces mitochondrial fragmentation and loss of network connectivity. Scale bars, 5 µm. Quantitative analyses of mitochondrial area, sphericity, and volume are shown for HEK293 cells (L–N) and COS7 cells (Q–S). Individual data points represent mitochondria; bars indicate mean ± SEM. For all panels, statistical significance is denoted as ns (not significant), p ≤ 0.05, p ≤ 0.01, *p ≤ 0.001, and **p ≤ 0.0001. Mitochondrial superoxide production was quantified using the MitoSOX™ Deep Red fluorescent probe and measured by a plate reader. Fluorescence intensity was normalized to control conditions. (T) Relative MitoSOX fluorescence following treatment with vehicle (DMSO), hydrogen peroxide (H₂O₂; 200 µM), and increasing doses of the indicated compound (30 µM and 50 µM), either alone or in combination with H₂O₂. (U) Independent experimental replicate showing consistent modulation of mitochondrial ROS across treatment conditions. H₂O₂ exposure significantly increased mitochondrial superoxide levels, whereas compound treatment reduced MitoSOX fluorescence in a dose-dependent manner. Co-treatment attenuated H₂O₂-induced mitochondrial ROS, with no significant differences observed between selected conditions as indicated (ns). Data are presented as mean ± SEM, with each dot representing an independent biological replicate. Statistical significance was assessed by one-way ANOVA with multiple-comparison correction; p < 0.05, p < 0.01, *p ≤ 0.001, and **p ≤ 0.0001; ns, not significant.

### Pharmacologic MICOS Inhibition Induces Conserved Mitochondrial Fragmentation and Alters Mitochondrial Redox Status

To determine whether acute pharmacologic disruption of MICOS recapitulates mitochondrial structural alterations observed upon genetic perturbation, we treated HEK293 cells with the MICOS inhibitor Miclixin (50 µM) and performed three-dimensional mitochondrial reconstruction. Vehicle-treated cells displayed interconnected mitochondrial networks with extensive branching (Figure 8J), whereas Miclixin-treated cells exhibited fragmented mitochondrial structures with reduced network connectivity (Figure 8K). Quantitative morphometric analysis revealed a significant reduction in mitochondrial area following MICOS inhibition compared with DMSO-treated controls (Figure 8L). In parallel, mitochondrial sphericity was significantly increased in Miclixin-treated cells (Figure 8M), indicating a shift toward more rounded mitochondrial morphology. Mitochondrial volume was also significantly reduced upon MICOS inhibition (Figure 8N), consistent with a loss of mitochondrial mass and network integrity.

To assess whether MICOS-dependent mitochondrial remodeling is conserved across cell types, we performed analogous experiments in COS7 cells treated with Miclixin (50 µM). Three-dimensional reconstructions showed that vehicle-treated COS7 cells maintained elongated, reticulated mitochondrial networks (Figure 8O), whereas MICOS inhibition led to pronounced mitochondrial fragmentation and reduced connectivity (Figure 8P). Quantitative analysis showed a significant decrease in mitochondrial area in Miclixin-treated COS7 cells compared with controls (Figure 8Q). Mitochondrial sphericity was significantly increased following MICOS inhibition (Figure 8R), indicating altered mitochondrial shape. In addition, mitochondrial volume was significantly reduced in Miclixin-treated cells compared with vehicle-treated controls (Figure 8S). These findings mirror the structural alterations observed in HEK293 cells, indicating a conserved requirement for MICOS integrity in maintaining mitochondrial morphology.

To examine the functional consequences of MICOS perturbation on mitochondrial redox status, mitochondrial superoxide levels were quantified using the MitoSOX™ Deep Red probe. Exposure to hydrogen peroxide (H₂O₂; 200 µM) significantly increased mitochondrial superoxide production relative to vehicle-treated controls (Figure 8T). Treatment with the indicated compound at 30 µM and 50 µM concentrations reduced MitoSOX fluorescence in a dose-dependent manner (Figure 8T). Co-treatment with H₂O₂ and the compound attenuated H₂O₂-induced mitochondrial superoxide levels, with no significant differences observed between selected treatment conditions as indicated (ns). An independent experimental replicate demonstrated consistent modulation of mitochondrial ROS across treatment groups (Figure 8U), confirming the reproducibility of the observed effects.

### Pharmacologic Disruption of Mitochondrial Inner Membrane Organization Co-ordinately Remodels Organelle Morphology and Inter-Organelle Interactions

To examine how perturbation of mitochondrial inner membrane organization influences cellular organelle architecture, we analyzed mitochondrial morphology and its relationship with lipid droplets and lysosomes following treatment with Myls22 or Miclxin. Three-dimensional confocal reconstructions revealed pronounced alterations in mitochondrial organisation compared with DMSO-treated controls (Figure 9A-C). Quantitative morphometric analysis demonstrated significant changes in mitochondrial area, volume, and sphericity upon treatment with either compound, indicating extensive remodeling of mitochondrial size and shape (Figure 9D-F).

**Figure 9.**
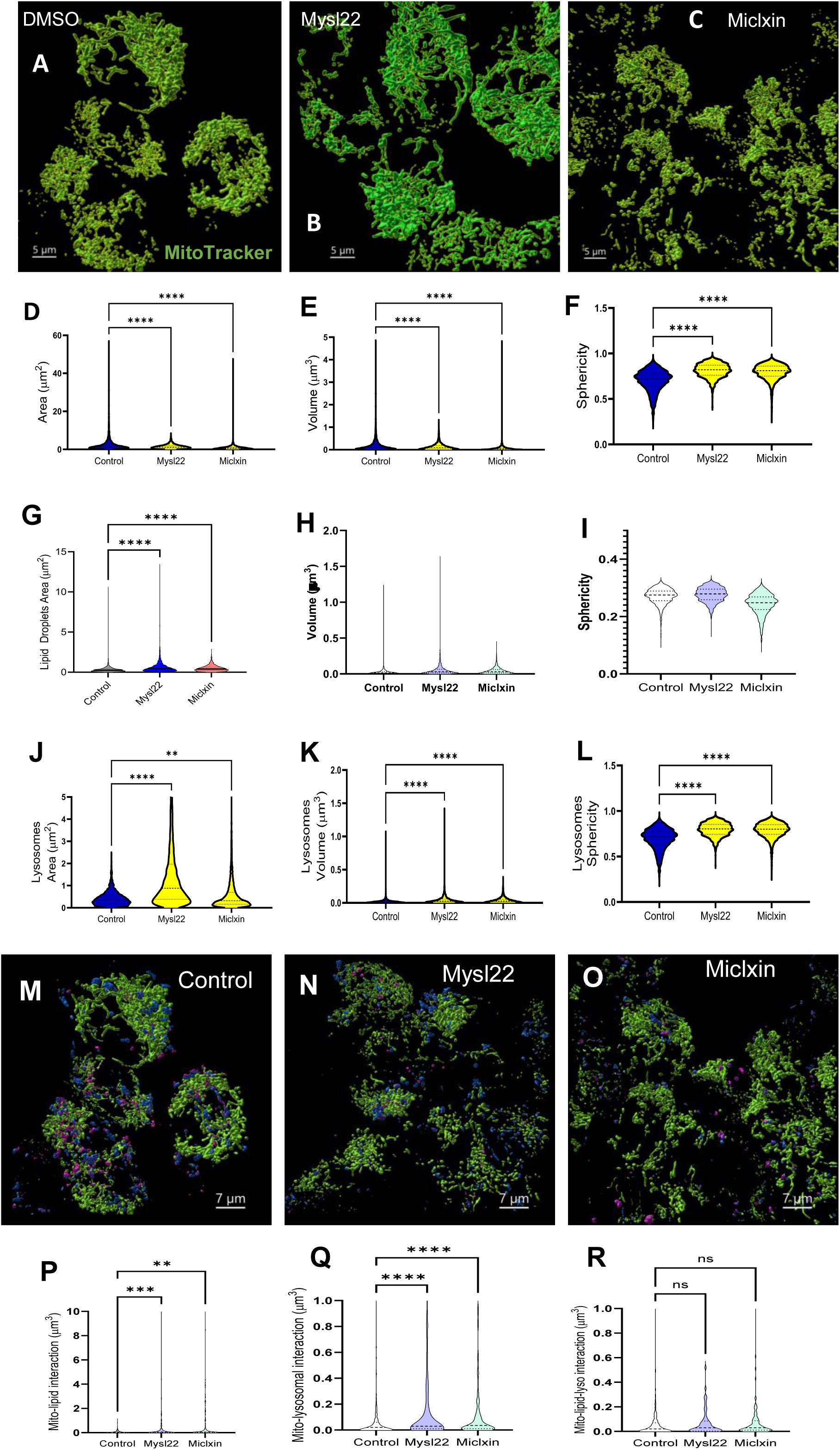
Pharmacological perturbation of mitochondrial inner membrane organization remodels mitochondrial, lipid droplet, and lysosomal morphology and inter-organelle interactions in HEK293 cells. (A–C) Representative 3D confocal reconstructions of mitochondria labeled with MitoTracker (green) in cells treated with DMSO (control) (A), the OPA1 inhibitor Mysl22 (B), or the MICOS complex inhibitor Miclxin (C). Scale bar, 5 μm. (D–F) Violin plot quantification of mitochondrial area (μm²) (D), volume (μm³) (E), and sphericity (F), revealing significant remodeling of mitochondrial size and shape following pharmacological inhibition of OPA1 or MICOS compared to control. (G–I) Lipid droplet morphometric analysis showing lipid droplet area (μm²) (G), volume (μm³) (H), and sphericity (I), indicating altered lipid storage and droplet morphology upon disruption of mitochondrial inner membrane organization. (J–L) Lysosomal morphometrics, including lysosome area (μm²) (J), volume (μm³) (K), and sphericity (L), demonstrate coordinated changes in lysosomal structure following Mysl22- or Miclxin-mediated perturbation. (M–O) Representative 3D reconstructions of mitochondria (green), lipid droplets (magenta), and lysosomes (blue) in control cells (M), following OPA1 inhibition with Mysl22 (N), or MICOS disruption with Miclxin (O). Scale bar, 7 μm. (P) Quantification of mitochondria–lipid droplet (mito–LD) interactions, showing a significant increase in organelle contacts following Mysl22 treatment relative to control. (Q) Quantification of mitochondria–lysosome (mito–lys) interactions, demonstrating enhanced contacts upon Mysl22 treatment, whereas Miclxin exhibits distinct or non-significant effects depending on the interaction type. (R) Quantification of three-way mitochondria–lipid droplet–lysosome (mito–LD–lys) interactions, defined as spatially coincident contacts in which a mitochondrion simultaneously engages both a lipid droplet and a lysosome within the same local volume. All violin plots represent pooled single-organelle measurements across cells from independent experiments. Statistical significance was assessed using one-way ANOVA with appropriate multiple-comparison correction; ns, not significant; **p < 0.01; ***p < 0.001; ****p < 0.0001.

Given the central role of mitochondria in lipid metabolism, we next assessed whether mitochondrial inner membrane perturbation was associated with changes in lipid droplet morphology. Both Myls22 and Miclxin treatments resulted in significant alterations in lipid droplet area, volume, and sphericity compared with control cells (Figure 9G-I), consistent with a reorganisation of cellular lipid storage properties in response to mitochondrial structural disruption. We further examined lysosomal morphology to determine whether mitochondrial inner membrane perturbation was accompanied by changes in the degradative compartment. Quantitative analysis revealed significant differences in lysosome area, volume, and sphericity following Myls22 and Miclxin treatment relative to controls (Figure 9J-L), suggesting coordinated remodeling of lysosomal structure alongside mitochondrial and lipid droplet alterations.

To investigate whether these morphological changes were accompanied by altered inter-organelle interactions, we performed three-dimensional interaction analyses between mitochondria, lipid droplets, and lysosomes. Representative reconstructions under control conditions and following OPA1 or MICOS perturbation illustrate distinct patterns of organelle proximity and organization (Figure 9 M-O). Quantitative analysis revealed a significant increase in mitochondria–lipid droplet interactions following Myls22 treatment compared to control cells (Figure 9P). Similarly, mitochondria–lysosome interactions were significantly enhanced upon Myls22 treatment, whereas Miclixin exhibited distinct or non-significant effects depending on the interaction type analysed (Figure 9Q). Finally, we assessed higher-order organelle organization by quantifying three-way mitochondria–lipid droplet–lysosome interactions. In contrast to the robust increases observed in pairwise interactions, three-way interactions did not exhibit a significant increase under mitochondrial inner membrane perturbation (Figure 9R), indicating that enhanced pairwise organelle contacts do not necessarily translate into stabilization of higher-order tri-organelle assemblies.

### Genetic disruption of OPA1 and MICOS components impairs mitochondrial respiration

#### Mouse kidney samples

To assess the impact of MICOS components and OPA1 on mitochondrial respiratory function *in vivo*, mitochondrial respiration was measured in mouse samples using the Seahorse XF Cell Mito Stress Test (Figure 10A). Basal oxygen consumption rate (OCR) was significantly reduced in samples from *Opa1* knockout (KO), *Chchd3* knockdown (KD), and *Chchd6* KD mice compared with controls (Figure 10B). Analysis of ATP-linked respiration revealed a significant decrease in *Opa1* KO and *Chchd6* KD samples, whereas *Chchd3* KD samples did not show a significant reduction relative to controls (Figure 10E). Maximal OCR was significantly reduced across *Opa1* KO, *Chchd3* KD, and *Chchd6* KD groups (Figure 10H), accompanied by a corresponding decrease in reserve respiratory capacity in these same conditions (Figure 10K). In parallel experiments targeting additional MICOS subunits, siRNA-mediated knockdown of *Mic10* or *Mic13* resulted in a significant reduction in basal OCR compared with control siRNA–treated samples (Figure 10C). ATP-linked respiration was modestly reduced following *Mic10* knockdown but not *Mic13* knockdown (Figure 10F). Maximal OCR and reserve capacity were both significantly reduced in *Mic10*- and *Mic13*-depleted samples relative to controls (Figure 10I, L).

**Figure 10.**
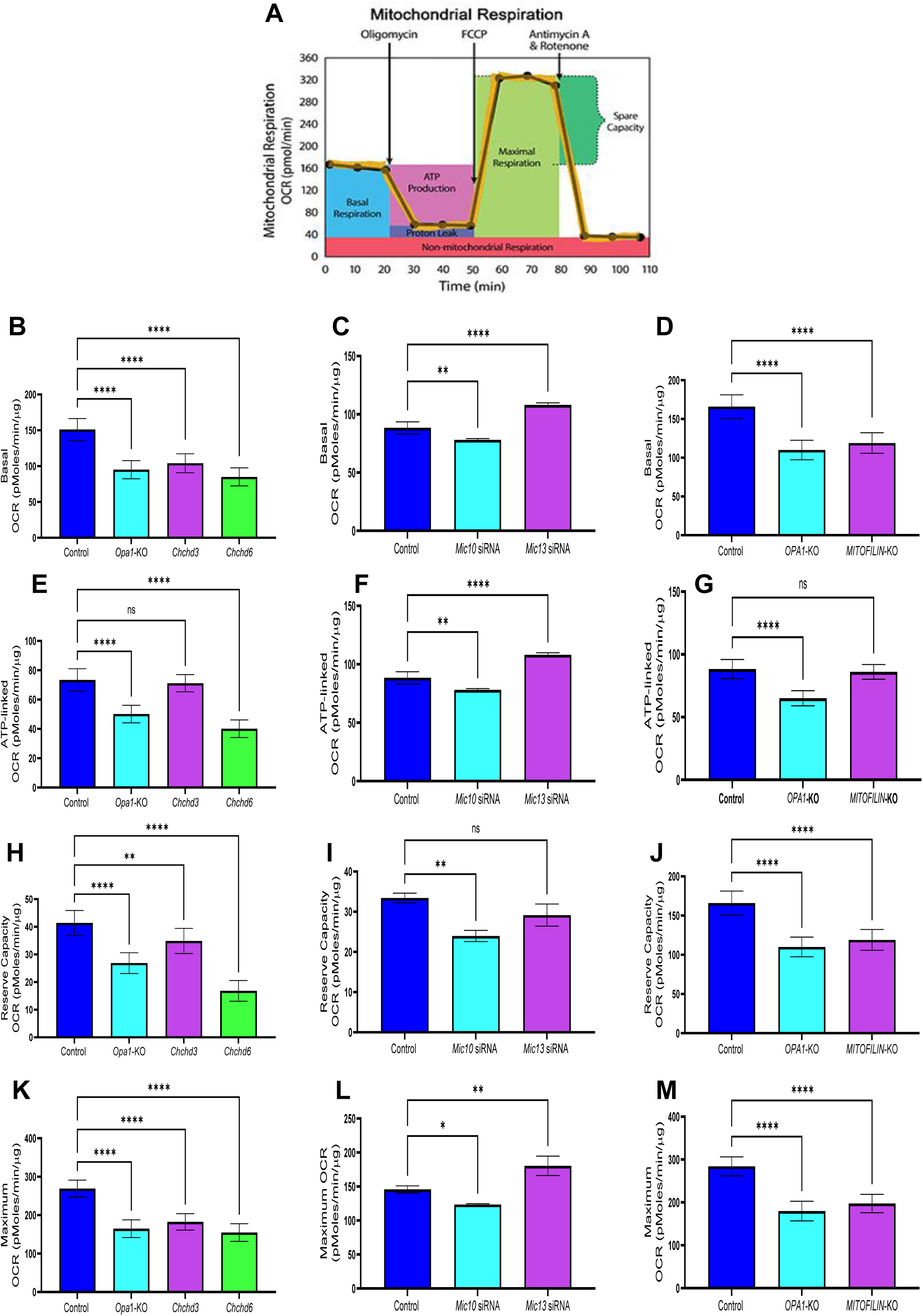
Genetic ablation of *Opa1* and siRNA-mediated knockdown of MICOS components impair mitochondrial respiration. (A) Schematic representation of the Agilent Seahorse XF Cell Mito Stress Test, illustrating changes in oxygen consumption rate (OCR) following sequential injection of oligomycin, FCCP, and antimycin A/rotenone, and the derived mitochondrial respiratory parameters, including basal respiration, ATP-linked respiration, maximal respiration, and reserve capacity. (B–M) Mitochondrial respiratory parameters measured using the Seahorse XF Cell Mito Stress Test. Panels are arranged vertically by experimental system and genetic perturbation. OCR values are normalized as indicated and presented as mean ± SEM. Each bar represents independent biological replicates. Statistical significance was determined using appropriate multiple-comparison analyses as indicated. Significance is denoted as ns (not significant), p ≤ 0.05, p ≤ 0.01, and **p ≤ 0.0001. **Mouse-derived data** (B, E, H, K) Basal OCR (B), ATP-linked OCR (E), maximal OCR (H), and reserve capacity OCR (K) were measured in mouse samples from control, *Opa1* knockout (KO), *Chchd3* siRNA–mediated knockdown (KD), and *Chchd6* siRNA–mediated knockdown (KD) groups. (C, F, I, L) Basal OCR (C), ATP-linked OCR (F), maximal OCR (I), and reserve capacity OCR (L) were measured in mouse samples treated with control siRNA, *Mic10* siRNA, or *Mic13* siRNA. **Cultured cell data** (D, G, J, M) Basal OCR (D), ATP-linked OCR (G), maximal OCR (J), and reserve capacity OCR (M) were measured in HEK293 cells under control conditions or following genetic ablation of OPA1 or MITOFILIN, as indicated.

#### HEK293 cell culture model

To determine whether MICOS-dependent respiratory defects are recapitulated in a cultured cell system, mitochondrial respiration was examined in HEK293 cells following genetic ablation of *OPA1* or Mitofilin. Basal OCR was significantly reduced in both *OPA1* KO and Mitofilin KO cells compared with control cells (Figure 10D). ATP-linked OCR was significantly decreased in *OPA1* KO cells but did not differ significantly between Mitofilin KO and control cells (Figure 10G). Maximal OCR was significantly reduced in both *OPA1* KO and Mitofilin KO cells relative to controls (Figure 10J). Similarly, reserve respiratory capacity was significantly diminished in both genetic perturbations compared with control cells (Figure 10M).

Across both mouse-derived samples and HEK293 cells, genetic disruption of OPA1 and multiple MICOS components consistently reduced basal and maximal mitochondrial respiration and diminished reserve respiratory capacity. However, ATP-linked respiration exhibited perturbation-specific effects that differed between individual MICOS components and between *in vivo* and *in vitro* systems.

### MICOS core components MIC60 and CHCHD6 are required for mitochondrial Ca²⁺ uptake, retention, and redox homeostasis

Mitochondrial Ca²⁺ handling is tightly linked to cristae architecture and inner membrane organization. To determine whether MICOS core components contribute directly to mitochondrial Ca²⁺ homeostasis, we examined mitochondrial Ca²⁺ uptake and retention in permeabilized HEK293 cells following siRNA-mediated depletion of MIC60 or CHCHD6. Efficient knockdown of both MICOS components was confirmed by immunoblotting, with ATP5A serving as a mitochondrial loading control (Figure 11A). Real-time measurements of mitochondrial Ca²⁺ uptake using the ratiometric indicator FuraFF revealed marked alterations in Ca²⁺ uptake kinetics upon MICOS depletion. Compared with scramble siRNA–treated cells, both MIC60- and CHCHD6-knockdown cells exhibited reduced mitochondrial Ca²⁺ uptake following a defined Ca²⁺ pulse, despite comparable experimental conditions and intact mitochondrial membrane potential prior to FCCP addition (Figure 11B). Quantitative analysis confirmed a significant reduction in Ca²⁺ uptake rates in both knockdown conditions relative to control cells (Figure 11C), indicating that MICOS integrity is required for efficient mitochondrial Ca²⁺ entry.

**Figure 11.**
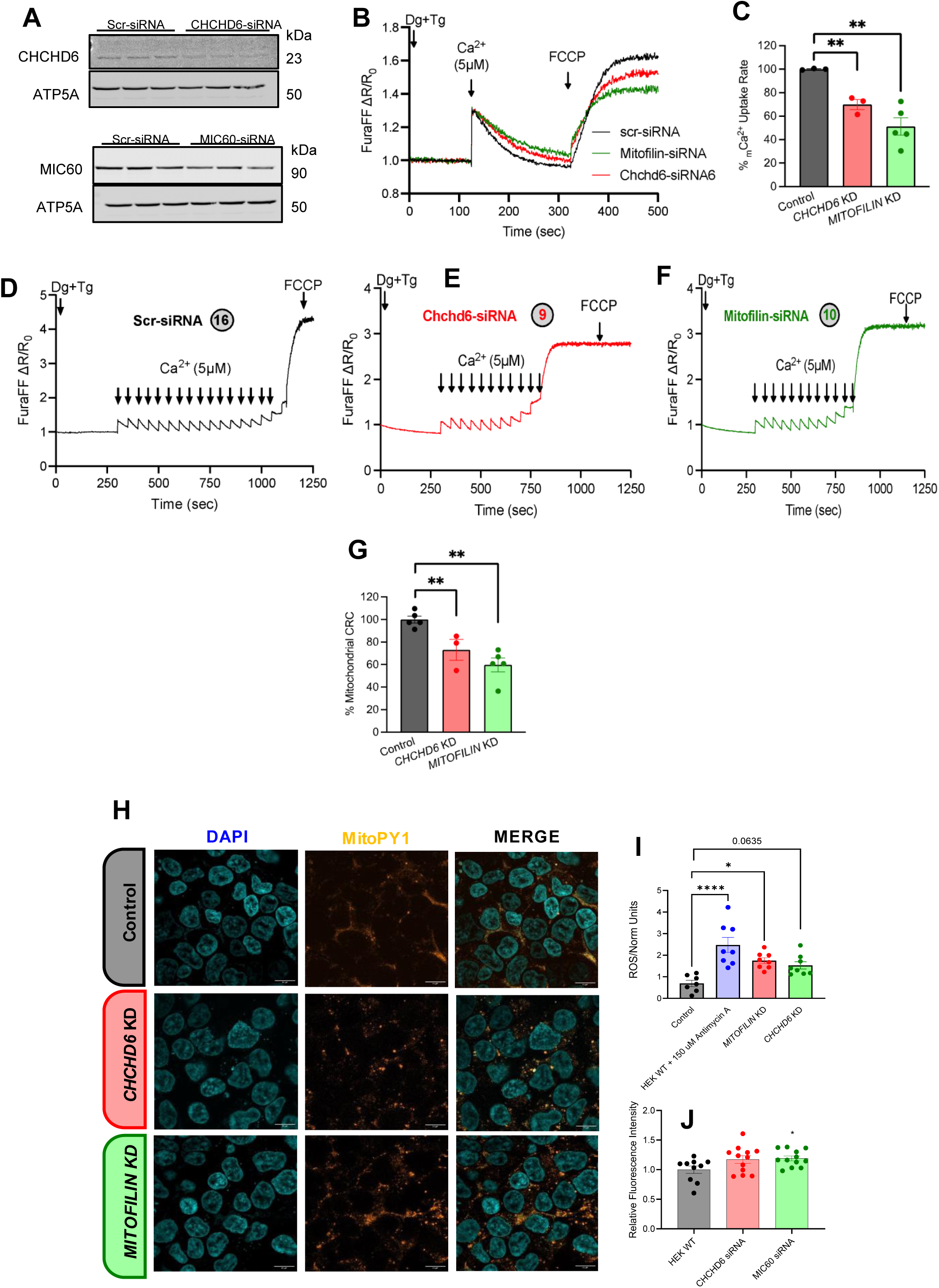
MICOS depletion disrupts mitochondrial calcium handling and redox balance in HEK293 cells. (A) Representative real-time traces of mitochondrial Ca²⁺ (mCa²⁺) uptake in permeabilized HEK293 cells transfected with scramble siRNA (scr-siRNA), MIC60-siRNA, or CHCHD6-siRNA. Traces were recorded using the ratiometric calcium indicator FuraFF, revealing altered mitochondrial calcium-uptake kinetics upon MICOS depletion. Digitonin (Dg) and thapsigargin (Tg) were added as indicated to permeabilize the plasma membrane and inhibit endoplasmic reticulum Ca²⁺ uptake, respectively, followed by a Ca²⁺ pulse (5 µM) and FCCP addition to collapse the mitochondrial membrane potential. (B) Immunoblot validation of siRNA-mediated knockdown efficiency in HEK293 cells. The top panel shows CHCHD6 protein levels following CHCHD6-siRNA transfection; the bottom panel shows MIC60 protein levels following MIC60-siRNA transfection. ATP5A was used as a mitochondrial loading control. (C–E) Representative mitochondrial calcium retention capacity (CRC) traces in permeabilized HEK293 cells transfected with scr-siRNA (C), CHCHD6-siRNA (D), or MIC60-siRNA (E). Sequential Ca²⁺ boluses (5 µM each; indicated by arrows) were administered until mitochondrial permeability transition pore opening, as evidenced by a sudden increase in cytosolic Ca²⁺. (F) Quantification of mitochondrial CRC calculated from traces shown in (C–E), expressed as a percentage relative to scr-siRNA control, demonstrating reduced calcium buffering capacity following MICOS component depletion. (G) Quantitative analysis of mitochondrial Ca²⁺ uptake rates derived from traces in (A), expressed as percentage change relative to control cells, revealing significant impairment upon MIC60 or CHCHD6 knockdown. Statistical analysis (panels A–G). For mitochondrial Ca²⁺ uptake kinetics and calcium retention capacity (CRC) analyses (A–G), representative traces are shown. Quantitative data in panels (F) and (G) were derived from independent biological replicates and are presented as mean ± SEM. Statistical significance was assessed using an unpaired two-tailed Student’s *t*-test or one-way ANOVA, as appropriate, with significance indicated as **P ≤ 0.01. (H) Representative fluorescence microscopy images of permeabilized HEK293 cells stained with DAPI (nuclei, blue) and the mitochondrial hydrogen peroxide sensor MitoPY1 (5 µM, 45 min at 37 °C), following transfection with scr-siRNA (control), MIC60-siRNA (Mitofilin KD), or CHCHD6-siRNA (CHCHD6 KD). Images were acquired at 60× magnification; merged channels are shown. (I) Quantification of mitochondrial ROS levels based on MitoPY1 fluorescence intensity from images shown in (H). (J) Relative MitoPY1 fluorescence emission intensity measured at 530 nm, normalized to control conditions, indicating altered mitochondrial redox status following MICOS depletion. Statistical analysis (panels H–J). For mitochondrial ROS measurements and MitoPY1 fluorescence quantification (H–J), data are presented as mean ± SEM, with each data point representing an independent biological replicate. Statistical significance was assessed using one-way ANOVA followed by Dunnett’s multiple-comparisons test. Significance levels are indicated as *P ≤ 0.05, **P ≤ 0.01, ***P ≤ 0.001, ****P ≤ 0.0001; ns, not significant.

We next assessed whether impaired Ca²⁺ uptake was accompanied by altered mitochondrial Ca²⁺ buffering capacity. Sequential Ca²⁺ bolus experiments demonstrated a pronounced reduction in mitochondrial calcium retention capacity (CRC) in MIC60- and CHCHD6-knockdown cells compared with controls (Figure 11D to F). In both knockdown conditions, mitochondria underwent permeability transition pore opening after fewer Ca²⁺ additions, indicating increased susceptibility to Ca²⁺-induced mitochondrial dysfunction. Quantification of CRC revealed a significant decrease in Ca²⁺ retention in MIC60- and CHCHD6-depleted cells relative to scramble siRNA–treated cells (Figure 11G).

Given the established coupling between mitochondrial Ca²⁺ dysregulation and oxidative stress, we next examined mitochondrial redox status following MICOS depletion. To assess mitochondrial redox status following MICOS depletion, mitochondrial hydrogen peroxide levels were measured in permeabilized HEK293 cells using the mitochondria-targeted H₂O₂ sensor MitoPY1. Representative fluorescence images revealed increased MitoPY1 signal intensity in cells transfected with MIC60-siRNA (Mitofilin KD) or CHCHD6-siRNA compared with scramble siRNA–treated controls (Figure 11H). Quantitative analysis of MitoPY1 fluorescence from microscopy images demonstrated a significant increase in mitochondrial ROS levels in both MIC60- and CHCHD6-depleted cells relative to control cells (Figure 11I). Antimycin A treatment served as a positive control, producing a robust increase in mitochondrial ROS signal. Consistent with microscopy-based measurements, quantification of MitoPY1 fluorescence emission intensity at 530 nm revealed a significant elevation in mitochondrial ROS levels following MIC60 or CHCHD6 knockdown compared with control conditions (Figure 11J).

Together, these data demonstrate that the MICOS core components MIC60 and CHCHD6 are essential for maintaining physiological mitochondrial Ca²⁺ uptake and retention. Disruption of MICOS integrity compromises mitochondrial Ca²⁺ handling, promotes early opening, and is associated with elevated mitochondrial oxidative stress, highlighting a critical role for cristae organization in safeguarding mitochondrial Ca²⁺ homeostasis.

## Discussion

### Structural Analysis

In the past, numerous studies have examined kidneys across various aging or disease states using TEM, which provides high-resolution 2D images^27,96,97^. While TEM is a useful technique for understanding changes in cristae integrity, it cannot accurately capture many structural details of mitochondria, such as the diverse structures they may adopt under cellular conditions^98^. We found that TEM area measurements directly contradict our findings of decreased mitochondrial volume with aging. Here, we utilized SBF-SEM to perform 3D reconstruction of the aged mouse kidney, which revealed many novel phenotypes and mitochondrial volume loss that were not otherwise captured by MRI or TEM. Using 3D reconstruction, we previously examined aged skeletal muscle in mice^37^. We observed increased mitochondrial fragmentation accompanied by a compensatory increase in its structural complexity. In aging murine kidneys, mitochondrial complexity did not undergo significant changes. We noted a high rate of diverse or unique mitochondrial shapes, which may, in turn, have functional implications.^29,98^ In this study, we found a large amount of variation in mitochondrial shape in both 3-month and 2-year old cohorts, which exhibit a mix of elongated, compact, large volume, small volume, nanotunnels, donut-shaped, and branching structural phenotypes. While our previous studies in skeletal and cardiac tissue showed a dominant phenotype across ageing, typically characterised by fragmentation, mitochondria in the kidney exhibit diverse shapes, and 3-month and 2-year samples do not show markedly different phenotypes.

One notable phenotype we observed is the presence of mitochondria in the form of a donut. We found that the 3-month samples displayed many branched forms of mitochondria with high complexity, and within them, donut-like structures formed. Past research utilizing 3D reconstruction in the aged brain of monkeys found a high rate of the donut mitochondria phenotype in the aged cohort, which had impaired memory function.^99^ Even beyond age, samples with mitochondrial donuts, which resulted in smaller synapses, had worse memory than older cohorts with normal mitochondria.^99^ While it is established that mitochondria donut are a hallmark of mitochondrial dysfunction^7^, interestingly, they seem to be differentially expressed in tissue types and correlate with each tissue’s functions. It has been suggested that their increased surface area relative to volume allows them to maintain more organelle contacts at the cost of lower ATP production^98^. It has also been found that, unlike swollen mitochondria, donut mitochondria maintain more of their internal structure and, potentially as a result, are not the target of mitophagy ^100^. Since our analysis shows that more donuts occur in the younger samples, these may represent a positive phenotype in some cases. Therefore, the exact roles of donut mitochondria still remain unclear and may extend beyond what has been previously hypothesized.

Beyond changes in relative bioenergetics between mitochondrial shapes, their roles in calcium homeostasis and other biomolecular pathways deserve further research. For instance, past studies have also suggested that decreasing the activity of the Akt pathway, which is upstream from the mTORC pathway, may be a mechanism to restore autophagy, clear out defective mitochondria, and restore biogenetics^101^. However, given that other studies have shown that donut mitochondria may not be subject to autophagic clearing of mitochondria^100^, suggesting a potential reason for different relative rates of mitochondrial donut. Additionally, research on mitochondrial aging in the kidney has found that mtDNA is more error-prone across aging, with up to a 5-fold increase in the number of point mutations and deletions^101^, which may also be responsible for the alterations in mitochondria structure as observed. Thus, the exact molecular underpinnings of these various shapes warrant further research.

Finally, we also found differences in mitochondrial structure across locations. Using 3D reconstruction, we found that in all ageing kidney samples, mitochondria are round and located next to the nucleus. The further away from the nucleus, the more unique mitochondrial structures are present, such as those with a large volume (increased mitochondrial function capacity), small volume, elongated shape (relatively greater surface area facilitates interaction with the surrounding environment), compact structure, nano tunnels, and donut-shaped appearance (increased surface area for interaction) are present^29^. The change in shape likely arises from mitochondrial stress ^98,102^. Whereas areas that are not undergoing stress likely present mitochondria that are typical and elongated. Therefore, different areas of the kidney may undergo stress, potentially linked to stress from filtration, while others are less susceptible to stress as aging progresses. In a previous study, we found that mitochondria in cardiac tissue retain their morphology^36^. Therefore, a similar mechanism may exist that helps retain morphology for intracellular regions, such as perinuclear kidney mitochondria. Notably, the kidney houses at least 16 types of epithelial cells^103^ and has distinct regions, including the cortex, medulla, and renal sections, which serve different functions^104^. However, our study did not permit the differentiation of these separate regions. Thus, future studies may consider using methods such as SDS-PAGE to further differentiate kidney samples^105^. Thus, future studies may further explore this by developing methods to separate kidney epithelial and globular areas using SBF-SEM and examining whether mitochondrial differences are region- or area-dependent across aging. Additionally, we were unable to perform sex comparisons, which may also reveal further heterogeneity across regions and between sexes.

### Structural–Metabolic Coupling Drives Redox Imbalance in the Aging Kidney

Across aging, we observed coordinated structural and metabolic remodeling consistent with oxidative stress arising, in part, from disruption of the MICOS complex. Oxidative stress activates poly(ADP-ribose) polymerase (PARP) as a DNA repair mechanism, which consumes nicotinamide adenine dinucleotide (NAD^+^), contributing to age-associated NAD^+^ decline in both males and females.^134^ Relevantly, extracellular NAD^+^ can trigger a cAMP-dependent pathway that promotes Ca^2+^ influx and generation of superoxide and nitric oxide.^135^ Reduced NAD^+^ availability has been implicated in mitochondrial dysfunction and diabetic kidney injury^136^, and studies using nicotinamide riboside to boost NAD^+^ levels suggest that simply restoring NAD^+^ pools may not fully rescue inter-organelle communication or cristae dynamics. ^137^ Together, these findings support the possibility that MICOS-dependent structural deterioration contributes to a redox environment that is not readily corrected by NAD^+^ repletion alone.

Our metabolic profiling revealed broad perturbations in amino acid, nucleotide, and redox metabolism in aged kidneys. We observed significant declines in methionine, valine, threonine, leucine, isoleucine, and glycine, highlighting disruption of pathways central to mitochondrial one-carbon metabolism and branched-chain amino acid utilization. Glycine, serine, and threonine metabolism—closely linked to the mitochondrial glycine cleavage system (GCS)—feed into purine biosynthesis and redox balance. Consistent with this, we detected marked reductions in purine intermediates, including inosine monophosphate (IMP), ribose-phosphate, and related nucleotide metabolites. Because the Pentose Phosphate Pathway (PPP) supplies ribose-5-phosphate and NADPH, impairment of this pathway may simultaneously compromise nucleotide synthesis and antioxidant capacity.^140^ PPP-derived NADPH is essential for glutathione (GSH) regeneration, and depletion of nucleotides such as xanthine and ribose promotes genomic instability.^141^

Interestingly, despite upstream perturbations in nucleotide biosynthesis, total adenylate pools (AMP, ADP, and ATP) were maintained in aged kidneys. This preservation of steady-state ATP suggests that energetic homeostasis may be buffered through compensatory mechanisms, even as redox and biosynthetic pathways are significantly remodeled. One interpretation is that aging kidneys prioritize ATP maintenance at the expense of redox flexibility and anabolic capacity. Alternatively, these findings may reflect a temporal hierarchy in which redox imbalance and nucleotide depletion precede overt energetic collapse.

Consistent with prior literature on dysregulated NAD^+^ metabolism in aging kidneys^142^, we observed significant declines in NAD^+^, NADP, and NAM, accompanied by accumulation of NADH (Figures 6H–K). This supports a redox shift hypothesis, wherein NAD^+^ is increasingly reduced to NADH without efficient re-oxidation.^143^ Such an imbalance is consistent with impaired electron transport chain (ETC) activity, particularly in the context of altered cristae architecture. Further reinforcing disruption of oxidative metabolism, we detected an age-dependent decline in FAD (Figure 6L). Similar to NAD(H), FAD functions as a critical electron carrier in oxidative phosphorylation, the TCA cycle, fatty acid β-oxidation, and ETC complex II activity. Together, reductions in NAD^+^, NADP, and FAD, along with NADH accumulation, point to impaired oxidative capacity and a shift toward reductive stress rather than simple ATP depletion.

Our lipidomic profiling revealed substantial age-associated remodeling of mitochondrial membrane lipids. TGOs and TGs serve as fatty acid storage pools and substrates for mitochondrial β-oxidation^144,145^, and their dysregulation may limit acetyl-CoA generation. Sterols contribute to mitochondrial membrane integrity and fluidity^146,147^, aligning with our structural observations of altered mitochondrial morphology. Of particular relevance, cardiolipins (DLCL and CL), phospholipids uniquely enriched in the inner mitochondrial membrane, are essential for cristae architecture and stabilization of respiratory chain supercomplexes.^152,153^Age-associated alterations in cardiolipin abundance and composition likely compromise ETC organization and electron flux. Such structural perturbations provide a plausible mechanistic link between membrane remodeling and the observed redox imbalance, including NADH accumulation and FAD depletion.

Additional lipid classes, including NAEs, LPIs, and Hex2Cer, also exhibited age-dependent changes. While NAEs are recognized mediators of endocannabinoid signaling^148,149^ and LPIs function in signaling pathways, their roles in kidney mitochondrial biology remain incompletely defined. Hex2Cer contributes to mitochondrial membrane composition^150,151^and may influence membrane curvature and organelle dynamics. Together, these lipidomic changes support a model in which age-dependent membrane remodeling destabilizes mitochondrial structure, impairs respiratory efficiency, and reinforces redox and metabolic rewiring.

Importantly, these metabolomic and lipidomic measurements were performed in whole-kidney tissue. While our structural analyses were specific to tubular cells, bulk tissue profiling may mask compartment- and cell-type–specific metabolic alterations, particularly within mitochondrial ATP pools. Therefore, although our data strongly associate mitochondrial structural remodeling with redox imbalance and metabolic reprogramming, additional compartment-resolved studies will be required to establish causality and define whether mitochondrial-localized ATP production is preserved despite systemic buffering. Collectively, our findings support a model in which age-associated disruption of MICOS-dependent mitochondrial architecture initiates a cascade of membrane remodeling, redox imbalance, impaired nucleotide biosynthesis, and compensatory energetic buffering. Rather than immediate ATP failure, aging kidneys exhibit a state of redox-constrained metabolism that may predispose them to injury under additional stress.

### The MICOS Complex as a Master Regulator

With respect to AKI and CKD, mitochondria are known to play a significant role in the pathophysiology of these diseases^106^. Mitochondrial dynamics are complex, and observing key regulators of mitochondrial form and function may explain the changes that occur in kidney disease states. Key regulators of mitochondria include *OPA1* (mitochondrial fusion) and *DRP1* (mitochondrial fission), which may be responsible for the changes observed in the kidney. Past research has shown that in AKI, *OPA1* expression decreases and *DRP1* expression increases, suggesting mitochondrial fragmentation ^2^. However, beyond models showing that decreased *DRP1* expression is not viable, mitochondrial fission is also important for maintaining various roles, including microtubule trafficking^107^. Therefore, this study sought to identify other targets and changes in mitochondrial structure beyond simple alterations in fusion and fission, which is often the extent of what TEM can survey. The MICOS complex is one such compelling target.

Aging in the kidneys is well-established by us and others to cause interstitial fibrosis and oxidative stress.^55,108–110^ Our results suggested that age-related loss of the MICOS complex leads to mitochondrial structural loss, generating oxidative stress and dysregulating calcium homeostasis. Since the MICOS complex forms across cristae junctions, the understanding of the interdependencies among MICOS complex proteins is still evolving. However, it is currently understood that some integral proteins, such as MIC60 (Mitofilin), regulate the expression of other proteins, including MIC10 and MIC19.^111^ Similarly, MIC60/MIC19 (Mitofilin/Chchd3), unlike other MICOS complex proteins, assemble independently of cardiolipin, with MIC19 being responsible for regulating the distribution of subcomplexes.^85^ Past studies of the MICOS complex in the kidney have been limited. However, they generally show that mitochondria-rich regions, including the kidney, have a high rate of MIC60 and its isoforms. A deletion of *Mitofilin* results in a lethal disruption of the overall complex.^111^ This underscores the central role of Mitofilin, relative to other components of the MICOS complex, with functions that extend beyond cristae and mitochondrial dynamics to nucleoid distribution, suggesting roles in mtDNA synthesis.^112^ This has been recapitulated by other studies, which show that Mitofilin depletion decreases mtDNA transcription, resulting in impaired bioenergetics in the kidney, as previously reviewed.^86^ Notably, we observed a marked age-related decrease in Mitofilin compared with other components; however, Mitofilin also showed a less drastic mitochondrial phenotype when knocked out than other MICOS complex proteins. While the structural analysis of MICOS complex knockouts is limited to TEM, this underscores the importance of considering other roles of the MICOS complex beyond its well-established, extensively reviewed function in cristae dynamics and biogenesis.^84,113^

The role of the MICOS complex in disease states remains controversial. Loss of the MICOS complex has been shown to reduce cardiac ATP levels, thereby impairing tissue integrity.^114^ Studies in other tissues, such as the liver, have shown that *CHCHD3* depletion impairs MERCs, leading to fatty liver disease with *SLC25A46* involvement.^115^ As previously reviewed, the MICOS complex has thus been involved in neurodegenerative disorders, metabolic syndromes, cardiac dysfunctions, and muscle pathologies^116^. In the kidney, as previously reviewed, impairment of Mitofilin has been specifically implicated in the pathophysiology of mtDNA-related renal diseases, diabetic kidney disease, kidney failure, and reperfusion.^86^ Interestingly, other studies have suggested a protective mechanism due to the loss of the MICOS complex. Loss of the MICOS complex, despite an aberrant cristae structure, is a protective factor against aging, as it exhibits an unexpected, pronounced lifespan extension in *Podospora anserina.*^117^ Specifically, it has been suggested that Miro-MIC60 interactions impair cellular respiration and cause oxidative stress, thereby preventing mitophagy and increasing susceptibility to Parkinson’s disease and Friedreich’s ataxia.^118^ This underscores the need to better understand the impact of the MICOS complex loss.

Notably, unlike other studies that show Miro-MIC60 interactions cause oxidative stress, we found that deletion of MICOS complex components Chchd6 and Mitofilin resulted in mitochondrial and cellular oxidative stress. As previously reviewed, oxidative stress has been observed following *Mitofilin* deletion in some tissues, such as the heart, but the interplay between the MICOS complex and oxidative stress remains poorly understood^86^. Notably, within the kidney, oxidative stress mediates age-associated renal cell death and has been linked to numerous pathological conditions, as previously reviewed.^93^ Since the loss of the MICOS complex is well understood to impair bioenergetics and ATP production, ^41,87^ our findings suggest that the closely linked process of free radical generation is also strengthened. MICOS-generated ROS may have various effects; for example, they can reduce NAD^+^, as observed in our aged tissue, leading to alterations in glycolysis, the TCA cycle, and oxidative phosphorylation, as previously reviewed.^119^ reviewed: changes in fuel availability alter TCA metabolite levels, with downstream effects on reduced mitochondrial calcium uptake and lowered matrix Ca^2+^ levels. , in turn, decreases Ca^2+^-dependent TCA cycle enzyme activity, including pyruvate dehydrogenase and α-ketoglutarate dehydrogenase, and, in some cases, induces autophagy as a compensatory mechanism for changes in substrate availability.^120^ Since we observed a concomitant decrease in mCa^2+^ uptake upon silencing of the MICOS complex, this suggests a vicious cycle in which ROS-dependent NAD+ and calcium-dependent TCA metabolites are lost due to MICOS; however, this pathway warrants further exploration. Alternatively, oxidative stress can cause mitochondrial permeability transition pore (mPTP) openings, which adaptively release excess ROS to maintain mitochondrial homeostasis; however, in pathological, persistent conditions, these pores can engage in destructive ROS-dependent ROS release^121,122^. While mPTP openings can be transient, calcium-dependent lowering of the membrane potential can also cause permanent openings, which confer an increased risk of apoptotic pathways^123^, suggesting an alternative pathway through which a feedback loop may arise due to ROS generation and calcium dysregulation following the silencing of proteins involved in the MICOS complex.

In murine renal tubular epithelial cells, an MCU-dependent increase in mitochondrial calcium accumulation leads to oxidative stress and, ultimately, senescence.^124^ This study examines the importance of further elucidating MICOS’s role in senescence and the therapies that target this process. For example, a recent study found that diminished Glis1 expression in age-related kidney aging models correlates with impaired mitochondrial quality control mechanisms. In contrast, increased Glis1 interaction with PGC-1α helps maintain mitochondrial stability, suggesting *Glis1* as a potential therapeutic target for mitigating cell senescence and age-related renal fibrosis.^125^ Furthermore, the role of the MICOS complex in regulating calcium highlights the importance of investigating other regulators of the mitochondrial Ca^2+^ uniporter (e.g., MICU1, MCU, EMRE), some of which have recently been shown to influence cristae morphology.^126^ While MICU1 has increasingly been shown to have a role in cristae morphology, the interconnectedness of these proteins has not yet been studied in the context of the MICOS complex’s downstream effectors^126,127^.

Accumulating evidence links nicotinamide adenine dinucleotide phosphate reduced oxidase (NOX)-driven oxidative stress to ER stress-induced apoptosis and subsequent renal dysfunction.^128^ Similarly, NOXs have been indicated to play a role in acute kidney injury by promoting oxidative stress.^33,129^ Beyond underscoring the therapeutic potential of NOXs, their interdependence with ER stress also highlights the importance of further studying MERCs. MERCs, which contact sites under 50 nm, that can be caused by ER stress, have previously been associated with calcium signaling and lipid metabolism. However, recent research has further suggested a potential role in senescence ^130^. Here, we did not comprehensively study MERCs, which are known to be implicated in mitochondrial calcium homeostasis.^131^ However, a qualitative analysis did show that wrappER forms principally in young samples. Past studies have shown that the rough endoplasmic reticulum may curve to closely wrap around the mitochondria and maintain lipid homeostasis, a phenomenon termed wrappER.^132^ Thus, the lipidomic shifts we observed with aging may be caused by deficient lipid flux and impaired cristae structure, without wrappER. This compartment, which performs numerous functions, including fatty acid secretion, may serve as an organelle linking mitochondria and peroxisomes to regulate overall lipid balance.^133^ Given that calcium homeostasis dysfunction is a potential avenue for kidney disease,^2^ it remains important to consider in the future how calcium homeostasis is impacted across aging through MERC modulation. Furthermore, these qualitative findings must be confirmed by an extensive quantitative study, similar to the one we performed for mitochondria.

## Conclusion

Our results demonstrate that the aging of murine kidney tissue is associated with cristae disarray and impaired mitochondrial structure, resulting in reduced organelle volume. This occurs alongside widespread metabolic and lipidomic shifts, as well as increased fibrosis and oxidative stress, which collectively reduce oxidative capacity and increase the risk of age-related disease states, including CKD and AKI.^38^ We further found that the MICOS complex is lost with kidney aging, in the absence of changes in other standard regulators of mitochondria and cristae morphology. While the age-dependent loss of the MICOS complex likely accounts for the loss of cristae architecture, silencing of MICOS components in HEK cells confers a structure similar to that of aged tissue. The MICOS complex silencing further causes oxidative stress. It is plausible that these changes result in a vicious cycle: MICOS loss drives oxidative stress, leading to calcium-dependent TCA dysregulation and NAD^+^ dysregulation, which in turn drive more oxidative stress and mtDNA loss, leading to a reduction in MICOS complex transcripts and a subsequent reduction in mitochondrial function, producing more oxidative stress byproducts, and ultimately leading to age-dependent disease states. Loss of the MICOS complex alters mitochondrial morphology, leading to distinct changes in mitochondrial structure. We frequently observed circular or ring-shaped mitochondria—commonly referred to as “donut” mitochondria—which are often associated with continuous mitochondrial constriction. The size of these mitochondrial dounts appears critical for modulating bending energy, a key biophysical barrier to their formation. This energy barrier may be counterbalanced by stress-inducible mitochondrial osmotic pressure. For instance, rotenone (a complex I inhibitor) and antimycin A (a complex III inhibitor) are known to induce reactive oxygen species (ROS), triggering a transition from tubular to donut - or blob-shaped mitochondria. These ROS-associated donut formations are typically transient and reversible, suggesting an adaptive response to oxidative stress.^29,98,154–156^ In our study, we also identified numerous compact mitochondria, further indicating disrupted mitochondrial dynamics. These morphological changes were consistently observed when comparing 3-month-old and 2-year-old tissue, with greater fragmentation in the aged samples. Transmission electron microscopy (TEM) confirmed these structural transitions. Together, these findings support a model in which the MICOS complex, together with OPA1, is essential for regulating mitochondrial dynamics and inner-membrane remodeling. This regulation maintains mitochondrial membrane potential, calcium buffering capacity, and cristae architecture—key components of mitochondrial function that are critical for proper kidney physiology. Interestingly, the ring- and donut -like mitochondrial structures observed following MICOS complex loss resemble those reported in our recent study of MFN-2–mediated mitochondrial remodeling in aged human skeletal muscle.^156^ In both contexts, we observed a consistent emergence of toroidal and compact mitochondrial forms, suggesting that despite acting on different structural components—the MICOS complex at the cristae junction and MFN-2 at the outer membrane tethering/fusion sites—both proteins influence mitochondrial architecture in a convergent manner. The buffering of mitochondrial membrane potential and the control of bending energy may represent shared mechanisms underlying these morphological transitions. Moreover, as we reported in Scudese et al., MFN-2 loss was associated with changes in mitochondrial volume, shape complexity, and network fragmentation, features also observed with MICOS deficiency. These findings suggest that inner and outer membrane remodeling pathways may be mechanistically linked through a shared biophysical constraint, such as osmotic stress or impaired fusion, that drives adaptive mitochondrial shaping under aging or stress conditions.

## DATA SHARING STATEMENT

Sharing of software, models, algorithms, protocols, methods, and other useful materials and resources related to the manuscript will be available on a public repository upon publication.

## FUNDING

All authors have no competing interests.

The BioVU project at VUMC is supported by numerous sources, including NIH-funded Shared Instrumentation Grants S10OD017985, S10RR025141, and S10OD025092, and CTSA grants UL1TR002243, UL1TR000445, and UL1RR024975. Its contents are solely the responsibility of the authors and do not necessarily represent official views of the National Centre for Advancing Translational Sciences. Genomic data are also supported by investigator-led projects, including U01HG004798, R01NS032830, RC2GM092618, P50GM115305, U01HG006378, U19HL065962, and R01HD074711, as well as additional funding sources listed at https://victr.vumc.org/biovu-funding/.

This project was funded by the National Institute of Health (NIH) NIDDK T-32, number DK007563 entitled Multidisciplinary Training in Molecular Endocrinology to Z.V.; National Institute of Health (NIH) NIDDK T-32, number DK007563 entitled Multidisciplinary Training in Molecular Endocrinology to A.C.; NSF MCB #2011577I to S.A.M.; The UNCF/Bristol-Myers Squibb E.E. Just Faculty Fund, Career Award at the Scientific Interface (CASI Award) from Burroughs Welcome Fund (BWF) ID # 1021868.01, BWF Ad-hoc Award, NIH Small Research Pilot Subaward to 5R25HL106365-12 from the National Institutes of Health PRIDE Program, DK020593, Vanderbilt Diabetes and Research Training Center for DRTC Alzheimer’s Disease Pilot & Feasibility Program. CZI Science Diversity Leadership grant number 2022- 253529 from the Chan Zuckerberg Initiative DAF, an advised fund of Silicon Valley Community Foundation, to A.H.J., and National Institutes of Health grant HD090061 and the Department of Veterans Affairs Office of Research Award I01 BX005352 to J.G. Howard Hughes Medical Institute Hanna H. Gray Fellows Program Faculty Phase (Grant# GT15655 awarded to M.R.M); and Burroughs Wellcome Fund PDEP Transition to Faculty (Grant# 1022604 awarded to M.R.M). National Institutes of Health Grants: R21DK119879 (to C.R.W.) and R01DK-133698 (to C.R.W.), American Heart Association Grant 16SDG27080009 (to C.R.W.) and by an American Society of Nephrology KidneyCure Transition to Independence Grant (to C.R.W.). Doris Duke Clinical Scientist Development Award grant 2021193, Burroughs Wellcome Fund grant 1021480, K23 HL156759, and R01 DK112262 (CNW). NIH Grants 2D43TW009744 (SKM), R21TW012635 (AK and SKM) and the American Heart Association Award Number 24IVPHA1297559 https://doi.org/10.58275/AHA.24IVPHA1297559.pc.gr.193866 (SKM). International federation for clinical chemistry (SKM). NIH Grants R01HL147818, R03HL155041, and R01HL144941 (A. Kirabo). NIH Grant R00DK120876 (D.T.), Harold S. Geneen Charitable Trust Awards Program (D.T.), Alzheimer’s Association AARG-NTF-23-1144888 (D.T.). NIH Grant R00AG065445 (P.J.), Alzheimer’s Association 24AARG-D-1191292 (P.J.), Wake ADRC REC and Development grant P30AG072947 (P.J.). American Heart Association Grant 23POST1020344 (A.K.). NIH K01AG062757 to (M.T.S.) ANRF (Anusandhan National Research Foundation), ANRF/ECRG/2024/001042/LS, ANRF/IRG/2024/001777/LS. IISER Tirupati, NFSG. (P.K) Confocal microscopy, super-resolution microscopy, and image processing were performed through the Vanderbilt Cell Imaging Shared Resource (CISR), supported by CA68485, DK58404, and EY08126. S10MH130456 funded the Nikon Spinning Disk Confocal SoRa microscope. Its contents are solely the responsibility of the authors and do not necessarily represent the official view of the NIH. The contents are solely the responsibility of the authors and do not necessarily represent the official view of the NIH. The funders had no role in study design, data collection, and analysis, decision to publish, or preparation of the manuscript. We would also like to acknowledge the Huck Institutes’ Metabolomics Core Facility (RRID:SCR_023864) for use of the OE 240 LCMS and Drs. Imhoi Koo, Ashley Shay, and Sergei Koshkin for helpful discussions on sample preparation and analysis.

**Conceptualization:** Antentor Hinton Jr., Melanie R. McReynolds

**Methodology:** Prasanna Katti, Praveena Prasad, Sepiso K. Masenga, Zer Vue, Prasanna Venkhatesh, Andrea G. Marshall, Benjamin Rodriguez, Han Le, Edgar Garza-Lopez, Alexandria Murphy, Brenita Jenkins, Ashlesha Kadam, Jianqiang Shao, Amber Crabtree, Pamela Martin, Chantell Evans, Mark A. Phillips, David Hubert, Nelson Wandira, Okwute M. Ochayi, Dhanendra Tomar, Clintoria R. Williams, Jennifer Gaddy, Briar Tomeau, LaCara Bell, Taneisha Gillyard, Markis’ Hamilton, Vineeta Sharma, Amadou Gaye, Elma Zaganjor, Olujimi A. Ajijola, Estevão Scudese, Tyne W. Miller Fleming, André Kinder, Chandravanu Dash, Anita M. Quintana, Bret C. Mobley, Julia D. Berry, Pooja Jadiya, Dao-Fu Dai, Annet Kirabo, Oleg Kovtun, Jenny C. Schafer, Sean Schaffer, Renata Oliveira Pereira

**Investigation:** Zer Vue, Praveena Prasad, Sepiso K. Masenga, Prasanna Katti, Prasanna Venkhatesh, Andrea G. Marshall, Benjamin Rodriguez, Han Le, Mohd M khan and all middle authors listed above.

**Formal Analysis:** Prasanna Katti, Zer Vue, Praveena Prasad, Sepiso K. Masenga, Prasanna Venkhatesh

**Visualisation:** Prasanna Katti, Zer Vue, Praveena Prasad, Sepiso K. Masenga, Prasanna Venkhatesh

**Writing – Original Draft:** Prasanna Katti, Zer Vue, Praveena Prasad, Sepiso K. Masenga, Mohd M Khan

**Writing – Review & Editing:** Prasanna Venkhatesh, Prasanna Katti, Sepiso K. Masenga, Mohd, M khan

**Resources:** Antentor Hinton Jr., Melanie R. McReynolds

**Supervision:** Antentor Hinton Jr., Melanie R. McReynolds

**Project Administration:** Antentor Hinton Jr., Melanie R. McReynolds

**Funding Acquisition:** Antentor Hinton Jr., Melanie R. McReynolds

## References

1. Ferguson, M. A. & Waikar, S. S. Established and Emerging Markers of Kidney Function. Clinical Chemistry 58, 680–689 (2012).

2. Ishimoto, Y. & Inagi, R. Mitochondria: a therapeutic target in acute kidney injury. Nephrology Dialysis Transplantation 31, 1062–1069 (2016).

3. Chronic Kidney Disease in the United States, 2019. Fluoride Action Network https://fluoridealert.org/studytracker/38332/ (2020).

4. Duann, P. & Lin, P.-H. Mitochondria Damage and Kidney Disease. in Mitochondrial Dynamics in Cardiovascular Medicine (ed. Santulli, G.) 529–551 (Springer International Publishing, Cham, 2017). doi:10.1007/978-3-319-55330-6_27.

5. Bhargava, P. & Schnellmann, R. G. Mitochondrial energetics in the kidney. Nat Rev Nephrol 13, 629–646 (2017).

6. Glancy, B. Visualizing mitochondrial form and function within the cell. Trends in molecular medicine 26, 58–70 (2020).

7. Picard, M. & McEwen, B. S. Mitochondria impact brain function and cognition. Proceedings of the National Academy of Sciences 111, 7–8 (2014).

8. Duchen, M. R. & Szabadkai, G. Roles of mitochondria in human disease. Essays in Biochemistry 47, 115–137 (2010).

9. Mao, J. et al. The relationship between kidney disease and mitochondria: a bibliometric study. Renal Failure 46, 2302963 (2024).

10. Parasyri, M. et al. Renal Phenotype in Mitochondrial Diseases: A Multicenter Study. Kidney Diseases 8, 148–159 (2022).

11. Forbes, J. M. & Thorburn, D. R. Mitochondrial dysfunction in diabetic kidney disease. Nat Rev Nephrol 14, 291–312 (2018).

12. Alway, S. E., Mohamed, J. S. & Myers, M. J. Mitochondria Initiate and Regulate Sarcopenia. Exerc Sport Sci Rev 45, 58–69 (2017).

13. Hepple, R. T. Mitochondrial Involvement and Impact in Aging Skeletal Muscle. Frontiers in Aging Neuroscience 6, (2014).

14. Flannery, P. J. & Trushina, E. Mitochondrial dynamics and transport in Alzheimer’s disease. Molecular and Cellular Neuroscience 98, 109–120 (2019).

15. Zhang, L. et al. Altered brain energetics induces mitochondrial fission arrest in Alzheimer’s Disease. Scientific reports 6, 1–12 (2016).

16. Boudina, S. et al. Mitochondrial energetics in the heart in obesity-related diabetes: direct evidence for increased uncoupled respiration and activation of uncoupling proteins. Diabetes 56, 2457–2466 (2007).

17. Friederich, M., Hansell, P. & Palm, F. Diabetes, oxidative stress, nitric oxide and mitochondria function. Current diabetes reviews 5, 120–144 (2009).

18. Venkatachalam, M. A. & Weinberg, J. M. The tubule pathology of septic acute kidney injury: a neglected area of research comes of age. Kidney International 81, 338–340 (2012).

19. Padovano, V., Podrini, C., Boletta, A. & Caplan, M. J. Metabolism and mitochondria in polycystic kidney disease research and therapy. Nat Rev Nephrol 14, 678–687 (2018).

20. Fieni, F., Bae Lee, S., Jan, Y. N. & Kirichok, Y. Activity of the mitochondrial calcium uniporter varies greatly between tissues. Nat Commun 3, 1317 (2012).

21. Figueiredo, P. A., Mota, M. P., Appell, H. J. & Duarte, J. A. The role of mitochondria in aging of skeletal muscle. Biogerontology 9, 67–84 (2008).

22. Lesnefsky, E. J., Chen, Q. & Hoppel, C. L. Mitochondrial Metabolism in Aging Heart. Circ Res 118, 1593–1611 (2016).

23. Zhang, J. et al. Alterations in mitochondrial dynamics with age-related Sirtuin1/Sirtuin3 deficiency impair cardiomyocyte contractility. Aging Cell 20, e13419 (2021).

24. Bratic, A. & Larsson, N.-G. The role of mitochondria in aging. The Journal of clinical investigation 123, 951–957 (2013).

25. Serviddio, G. et al. Bioenergetics in aging: mitochondrial proton leak in aging rat liver, kidney and heart. Redox Report 12, 91–95 (2007).

26. Yamamoto, T. et al. Time-dependent dysregulation of autophagy: Implications in aging and mitochondrial homeostasis in the kidney proximal tubule. Autophagy 12, 801–813 (2016).

27. Cui, J. et al. Age-related changes in the function of autophagy in rat kidneys. AGE 34, 329–339 (2012).

28. Neikirk, K. et al. Call to Action to Properly Utilize Electron Microscopy to Measure Organelles to Monitor Disease. European Journal of Cell Biology 151365 (2023) doi:10.1016/j.ejcb.2023.151365.

29. Jenkins, B. C. et al. Mitochondria in disease: changes in shapes and dynamics. Trends Biochem Sci 49, 346–360 (2024).

30. Marshall, A. G. et al. Serial Block Face-Scanning Electron Microscopy as a Burgeoning Technology. Adv Biol (Weinh) e2300139 (2023) doi:10.1002/adbi.202300139.

31. Courson, J. A. et al. Serial Block-Face Scanning Electron Microscopy (SBF-SEM) of Biological Tissue Samples. J Vis Exp 10.3791/62045 (2021) doi:10.3791/62045.

32. Dutta, S. & Sengupta, P. Men and mice: Relating their ages. Life Sci 152, 244–248 (2016).

33. Li, M. S. et al. NADPH oxidase-2 mediates zinc deficiency-induced oxidative stress and kidney damage. American Journal of Physiology-Cell Physiology 312, C47–C55 (2017).

34. Tomsa, A. M., Alexa, A. L., Junie, M. L., Rachisan, A. L. & Ciumarnean, L. Oxidative stress as a potential target in acute kidney injury. PeerJ 7, e8046 (2019).

35. Genin, E. C. et al. CHCHD 10 mutations promote loss of mitochondrial cristae junctions with impaired mitochondrial genome maintenance and inhibition of apoptosis. EMBO molecular medicine 8, 58–72 (2016).

36. Vue, Z. et al. Three-Dimensional Mitochondria Reconstructions of Murine Cardiac Muscle Changes in Size Across Aging. American Journal of Physiology-Heart and Circulatory Physiology (2023) doi:10.1152/ajpheart.00202.2023.

37. Vue, Z. et al. Mouse Skeletal Muscle Decrease in the MICOS Complex and Altered Mitochondrial Networks with age. bioRxiv 2022–03 (2022).

38. Gyurászová, M., Gurecká, R., Bábíčková, J. & Tóthová, Ľ. Oxidative Stress in the Pathophysiology of Kidney Disease: Implications for Noninvasive Monitoring and Identification of Biomarkers. Oxid Med Cell Longev 2020, 5478708 (2020).

39. Hill Gallant, K. M. & Spiegel, D. M. Calcium Balance in Chronic Kidney Disease. Curr Osteoporos Rep 15, 214–221 (2017).

40. Pereira, R. O. et al. OPA 1 deficiency promotes secretion of FGF 21 from muscle that prevents obesity and insulin resistance. The EMBO journal 36, 2126–2145 (2017).

41. Vue, Z. et al. 3D reconstruction of murine mitochondria reveals changes in structure during aging linked to the MICOS complex. Aging Cell 22, e14009 (2023).

42. Danciu, I. et al. Secondary use of clinical data: the Vanderbilt approach. J Biomed Inform 52, 28–35 (2014).

43. Roden, D. M. et al. Development of a large-scale de-identified DNA biobank to enable personalized medicine. Clin Pharmacol Ther 84, 362–369 (2008).

44. Dennis, J. K. et al. Clinical laboratory test-wide association scan of polygenic scores identifies biomarkers of complex disease. Genome Medicine 13, 6 (2021).

45. 1000 Genomes Project Consortium et al. A global reference for human genetic variation. Nature 526, 68–74 (2015).

46. Price, A. L. et al. Principal components analysis corrects for stratification in genome-wide association studies. Nat Genet 38, 904–909 (2006).

47. Das, S. et al. Next-generation genotype imputation service and methods. Nat Genet 48, 1284–1287 (2016).

48. McCarthy, S. et al. A reference panel of 64,976 haplotypes for genotype imputation. Nat Genet 48, 1279–1283 (2016).

49. The GTEx Consortium atlas of genetic regulatory effects across human tissues. Science 369, 1318–1330 (2020).

50. Gamazon, E. R. et al. A gene-based association method for mapping traits using reference transcriptome data. Nat Genet 47, 1091–1098 (2015).

51. Hu, Y. et al. A statistical framework for cross-tissue transcriptome-wide association analysis. Nat Genet 51, 568–576 (2019).

52. Zhou, D. et al. A unified framework for joint-tissue transcriptome-wide association and Mendelian randomization analysis. Nat Genet 52, 1239–1246 (2020).

53. Carroll, R. J., Bastarache, L. & Denny, J. C. R PheWAS: data analysis and plotting tools for phenome-wide association studies in the R environment. Bioinformatics 30, 2375–2376 (2014).

54. Wu, P. et al. Mapping ICD-10 and ICD-10-CM Codes to Phecodes: Workflow Development and Initial Evaluation. JMIR Med Inform 7, e14325 (2019).

55. Chu, Y. et al. Glutathione peroxidase-1 overexpression reduces oxidative stress, and improves pathology and proteome remodeling in the kidneys of old mice. Aging Cell 19, e13154 (2020).

56. Garza-Lopez, E. et al. Protocols for Generating Surfaces and Measuring 3D Organelle Morphology Using Amira. Cells 11, 65 (2022).

57. Neikirk, K. et al. Systematic Transmission Electron Microscopy-Based Identification and 3D Reconstruction of Cellular Degradation Machinery. Advanced Biology 7, 2200221 (2023).

58. Hinton, A. et al. A Comprehensive Approach to Sample Preparation for Electron Microscopy and the Assessment of Mitochondrial Morphology in Tissue and Cultured Cells. Adv Biol (Weinh) e2200202 (2023) doi:10.1002/adbi.202200202.

59. Lu, W., Wang, L., Chen, L., Hui, S. & Rabinowitz, J. D. Extraction and Quantitation of Nicotinamide Adenine Dinucleotide Redox Cofactors. Antioxidants & Redox Signaling 28, 167–179 (2018).

60. Wang, L. et al. Peak Annotation and Verification Engine for Untargeted LC–MS Metabolomics. Anal. Chem. 91, 1838–1846 (2019).

61. Adusumilli, R. & Mallick, P. Data Conversion with ProteoWizard msConvert. in Proteomics: Methods and Protocols (eds. Comai, L., Katz, J. E. & Mallick, P.) 339–368 (Springer, New York, NY, 2017). doi:10.1007/978-1-4939-6747-6_23.

62. Lam, J. et al. A Universal Approach to Analyzing Transmission Electron Microscopy with ImageJ. Cells 10, 2177 (2021).

63. Hinton, A. et al. A comprehensive approach for artifact-free sample preparation and assessment of mitochondrial morphology in tissue and cultured cells. bioRxiv (2021).

64. Childs, D. D. et al. In-phase signal intensity loss in solid renal masses on dual-echo gradient-echo MRI: association with malignancy and pathologic classification. AJR Am J Roentgenol 203, W421–428 (2014).

65. Outwater, E. K., Bhatia, M., Siegelman, E. S., Burke, M. A. & Mitchell, D. G. Lipid in renal clear cell carcinoma: detection on opposed-phase gradient-echo MR images. Radiology 205, 103–107 (1997).

66. Fang, Y. et al. The ageing kidney: Molecular mechanisms and clinical implications. Ageing Research Reviews 63, 101151 (2020).

67. Wang, K., Liao, Q. & Chen, X. Research progress on the mechanism of renal interstitial fibrosis in obstructive nephropathy. Heliyon 9, e18723 (2023).

68. Menn-Josephy, H. et al. Renal interstitial fibrosis: an imperfect predictor of kidney disease progression in some patient cohorts. Am J Nephrol 44, 289–299 (2016).

69. Bandookwala, M. & Sengupta, P. 3-Nitrotyrosine: a versatile oxidative stress biomarker for major neurodegenerative diseases. Int J Neurosci 130, 1047–1062 (2020).

70. Lv, W., Booz, G. W., Fan, F., Wang, Y. & Roman, R. J. Oxidative Stress and Renal Fibrosis: Recent Insights for the Development of Novel Therapeutic Strategies. Front Physiol 9, 105 (2018).

71. Walker, L. M. et al. Oxidative Stress and Reactive Nitrogen Species Generation during Renal Ischemia. Toxicological Sciences 63, 143–148 (2001).

72. Qian, J. et al. Nitrotyrosine Level Was Associated with Mortality in Patients with Acute Kidney Injury. PLOS ONE 8, e79962 (2013).

73. Brandt, T. et al. Changes of mitochondrial ultrastructure and function during ageing in mice and Drosophila. eLife 6, e24662.

74. L, G., Nn, van der W., Ij, O., Pj, P. & Am, van der B. Loss of the intermembrane space protein Mgm1/OPA1 induces swelling and localized constrictions along the lengths of mitochondria. The Journal of biological chemistry 279, (2004).

75. Cogliati, S., Enriquez, J. A. & Scorrano, L. Mitochondrial cristae: where beauty meets functionality. Trends in biochemical sciences 41, 261–273 (2016).

76. Eisner, V. et al. Mitochondrial fusion dynamics is robust in the heart and depends on calcium oscillations and contractile activity. Proc Natl Acad Sci U S A 114, E859–E868 (2017).

77. Crabtree, A. et al. Defining Mitochondrial Cristae Morphology Changes Induced by Aging in Brown Adipose Tissue. Adv Biol (Weinh) e2300186 (2023) doi:10.1002/adbi.202300186.

78. Ilacqua, N., Anastasia, I. & Pellegrini, L. Isolation and analysis of fractions enriched in WrappER-associated mitochondria from mouse liver. STAR Protocols 2, 100752 (2021).

79. Lombardi, G. et al. Sex differences in chronic kidney disease–related complications and mortality across levels of glomerular filtration rate. Nephrology Dialysis Transplantation gfae087 (2024) doi:10.1093/ndt/gfae087.

80. Melsom, T. et al. Sex Differences in Age-Related Loss of Kidney Function. Journal of the American Society of Nephrology : JASN 33, 1891 (2022).

81. Ansari, A., Walton, S. L. & Denton, K. M. Sex- and age-related differences in renal and cardiac injury and senescence in stroke-prone spontaneously hypertensive rats. Biol Sex Differ 14, 1–14 (2023).

82. McBride, E. L. et al. Comparison of 3D cellular imaging techniques based on scanned electron probes: Serial block face SEM vs. Axial bright-field STEM tomography. Journal of Structural Biology 202, 216–228 (2018).

83. Li, Q. & Hoppe, T. Role of amino acid metabolism in mitochondrial homeostasis. Front Cell Dev Biol 11, 1127618 (2023).

84. Anand, R., Reichert, A. S. & Kondadi, A. K. Emerging Roles of the MICOS Complex in Cristae Dynamics and Biogenesis. Biology (Basel) 10, 600 (2021).

85. Friedman, J. R., Mourier, A., Yamada, J., McCaffery, J. M. & Nunnari, J. MICOS coordinates with respiratory complexes and lipids to establish mitochondrial inner membrane architecture. Elife 4, e07739 (2015).

86. Feng, Y., Madungwe, N. B. & Bopassa, J. C. Mitochondrial inner membrane protein, Mic60/mitofilin in mammalian organ protection. J Cell Physiol 234, 3383–3393 (2019).

87. An, J. et al. CHCM1/CHCHD6, Novel Mitochondrial Protein Linked to Regulation of Mitofilin and Mitochondrial Cristae Morphology *. Journal of Biological Chemistry 287, 7411–7426 (2012).

88. Darshi, M. et al. ChChd3, an Inner Mitochondrial Membrane Protein, Is Essential for Maintaining Crista Integrity and Mitochondrial Function *. Journal of Biological Chemistry 286, 2918–2932 (2011).

89. Darshi, M. & Taylor, S. S. Mitochondrial ChChD3 acts as a Scaffold for Mitofilin, Sam50 and PKA. (2008).

90. Barrera, M., Koob, S., Dikov, D., Vogel, F. & Reichert, A. S. OPA1 functionally interacts with MIC60 but is dispensable for crista junction formation. FEBS Letters 590, 3309–3322 (2016).

91. Hu, C. et al. OPA1 and MICOS Regulate mitochondrial crista dynamics and formation. Cell Death Dis 11, 1–17 (2020).

92. Gilkerson, R., De La Torre, P. & St. Vallier, S. Mitochondrial OMA1 and OPA1 as Gatekeepers of Organellar Structure/Function and Cellular Stress Response. Frontiers in Cell and Developmental Biology 9, (2021).

93. Percy, C., Pat, B., Poronnik, P. & Gobe, G. Role of oxidative stress in age-associated chronic kidney pathologies. Advances in Chronic Kidney Disease 12, 78–83 (2005).

94. Stenvinkel, P. et al. Chronic Inflammation in Chronic Kidney Disease Progression: Role of Nrf2. Kidney Int Rep 6, 1775–1787 (2021).

95. Ding, C. et al. Mitofilin and CHCHD6 physically interact with Sam50 to sustain cristae structure. Scientific reports 5, 1–11 (2015).

96. Bárcena, C., Martínez, M. A., Ortega, M. P., Muñoz, H. G. & Sárraga, G. U. Mitochondria with Tubulovesicular Cristae in Renal Oncocytomas. Ultrastructural Pathology 34, 315–320 (2010).

97. O’Toole, J. F., Patel, H. V., Naples, C. J., Fujioka, H. & Hoppel, C. L. Decreased cytochrome c mediates an age-related decline of oxidative phosphorylation in rat kidney mitochondria. Biochemical Journal 427, 105–112 (2010).

98. Glancy, B., Kim, Y., Katti, P. & Willingham, T. B. The Functional Impact of Mitochondrial Structure Across Subcellular Scales. Frontiers in Physiology 11, (2020).

99. Hara, Y. et al. Presynaptic mitochondrial morphology in monkey prefrontal cortex correlates with working memory and is improved with estrogen treatment. Proceedings of the National Academy of Sciences 111, 486–491 (2014).

100. Zhou, Y. et al. Topology-dependent, bifurcated mitochondrial quality control under starvation. Autophagy 16, 562–574 (2020).

101. Jankauskas, S. S. et al. Aged kidney: can we protect it? Autophagy, mitochondria and mechanisms of ischemic preconditioning. Cell Cycle 17, 1291–1309 (2018).

102. Long, Q. et al. Modeling of Mitochondrial Donut Formation. Biophysical journal 109, 892–9 (2015).

103. Balzer, M. S., Rohacs, T. & Susztak, K. How Many Cell Types Are in the Kidney and What Do They Do? Annu Rev Physiol 84, 507–531 (2022).

104. Agarwal, S. K., Sethi, S. & Dinda, A. K. Basics of kidney biopsy: A nephrologist’s perspective. Indian J Nephrol 23, 243–252 (2013).

105. Samuel, C. S. Determination of Collagen Content, Concentration, and Sub-types in Kidney Tissue. in Kidney Research: Experimental Protocols (eds. Becker, G. J. & Hewitson, T. D.) 223–235 (Humana Press, Totowa, NJ, 2009). doi:10.1007/978-1-59745-352-3_16.

106. Jiang, M. et al. Mitochondrial dysfunction and the AKI-to-CKD transition. American Journal of Physiology-Renal Physiology 319, F1105–F1116 (2020).

107. Bowes, T. & Gupta, R. S. Novel mitochondrial extensions provide evidence for a link between microtubule-directed movement and mitochondrial fission. Biochemical and Biophysical Research Communications 376, 40–45 (2008).

108. Gomes, P. et al. Aging increases oxidative stress and renal expression of oxidant and antioxidant enzymes that are associated with an increased trend in systolic blood pressure. Oxid Med Cell Longev 2, 138–145 (2009).

109. Zhou, X. J. et al. The aging kidney. Kidney International 74, 710–720 (2008).

110. O’Sullivan, E. D., Hughes, J. & Ferenbach, D. A. Renal Aging: Causes and Consequences. J Am Soc Nephrol 28, 407–420 (2017).

111. Rockfield, S. M. et al. Genetic ablation of Immt induces a lethal disruption of the MICOS complex. Life Sci Alliance 7, e202302329 (2024).

112. Li, H. et al. Mic60/Mitofilin determines MICOS assembly essential for mitochondrial dynamics and mtDNA nucleoid organization. Cell Death & Differentiation 23, 380–392 (2016).

113. Viana, M. P., Levytskyy, R. M., Anand, R., Reichert, A. S. & Khalimonchuk, O. Protease OMA1 modulates mitochondrial bioenergetics and ultrastructure through dynamic association with MICOS complex. iScience 24, 102119 (2021).

114. Birker, K. et al. Mitochondrial MICOS complex genes, implicated in hypoplastic left heart syndrome, maintain cardiac contractility and actomyosin integrity. eLife 12, e83385 (2023).

115. Dong, J. et al. Mic19 depletion impairs endoplasmic reticulum-mitochondrial contacts and mitochondrial lipid metabolism and triggers liver disease. Nat Commun 15, 168 (2024).

116. Eramo, M. J., Lisnyak, V., Formosa, L. E. & Ryan, M. T. The ‘mitochondrial contact site and cristae organising system’ (MICOS) in health and human disease. The Journal of Biochemistry 167, 243–255 (2020).

117. Warnsmann, V. et al. Disruption of the MICOS complex leads to an aberrant cristae structure and an unexpected, pronounced lifespan extension in Podospora anserina. Journal of Cellular Biochemistry 123, 1306–1326 (2022).

118. Li, L. et al. A mitochondrial membrane-bridging machinery mediates signal transduction of intramitochondrial oxidation. Nat Metab 3, 1242–1258 (2021).

119. Yang, Y. & Sauve, A. A. NAD+ metabolism: Bioenergetics, signaling and manipulation for therapy. Biochim Biophys Acta 1864, 1787–1800 (2016).

120. Tomar, D. & Elrod, J. W. Metabolite regulation of the mitochondrial calcium uniporter channel. Cell Calcium 92, 102288 (2020).

121. Zorov, D. B., Juhaszova, M. & Sollott, S. J. Mitochondrial Reactive Oxygen Species (ROS) and ROS-Induced ROS Release. Physiol Rev 94, 909–950 (2014).

122. Batandier, C., Leverve, X. & Fontaine, E. Opening of the mitochondrial permeability transition pore induces reactive oxygen species production at the level of the respiratory chain complex I. J Biol Chem 279, 17197–17204 (2004).

123. Vianello, A. et al. The mitochondrial permeability transition pore (PTP) — An example of multiple molecular exaptation? Biochimica et Biophysica Acta (BBA) - Bioenergetics 1817, 2072–2086 (2012).

124. Xiong, Y. et al. Mitochondrial calcium uniporter promotes kidney aging in mice through inducing mitochondrial calcium-mediated renal tubular cell senescence. Acta Pharmacol Sin 1–14 (2024) doi:10.1038/s41401-024-01298-5.

125. Xu, L. et al. GLIS1 alleviates cell senescence and renal fibrosis through PGC1-α mediated mitochondrial quality control in kidney aging. Free Radical Biology and Medicine 209, 171–184 (2023).

126. Gottschalk, B. et al. MICU1 controls cristae junction and spatially anchors mitochondrial Ca2+ uniporter complex. Nat Commun 10, 3732 (2019).

127. Tomar, D. et al. MICU1 regulates mitochondrial cristae structure and function independently of the mitochondrial Ca2+ uniporter channel. Sci Signal 16, eabi8948 (2023).

128. Li, G., Scull, C., Ozcan, L. & Tabas, I. NADPH oxidase links endoplasmic reticulum stress, oxidative stress, and PKR activation to induce apoptosis. Journal of Cell Biology 191, 1113–1125 (2010).

129. Jeong, B. Y. et al. Oxidative stress caused by activation of NADPH oxidase 4 promotes contrast-induced acute kidney injury. PLOS ONE 13, e0191034 (2018).

130. Ziegler, D. V., Martin, N. & Bernard, D. Cellular senescence links mitochondria-ER contacts and aging. Commun Biol 4, 1–14 (2021).

131. Carreras-Sureda, A. et al. Non-canonical function of IRE1α determines mitochondria-associated endoplasmic reticulum composition to control calcium transfer and bioenergetics. Nat Cell Biol 21, 755–767 (2019).

132. Anastasia, I. et al. Mitochondria-rough-ER contacts in the liver regulate systemic lipid homeostasis. Cell Reports 34, 108873 (2021).

133. Ilacqua, N. et al. A three-organelle complex made by wrappER contacts with peroxisomes and mitochondria responds to liver lipid flux changes. Journal of Cell Science 135, jcs259091 (2021).

134. Massudi, H., et al. Age-Associated Changes In Oxidative Stress and NAD+ Metabolism In Human Tissue. PLOS ONE 7, e42357 (2012).

135. Bruzzone, S. et al. Extracellular NAD+ regulates intracellular calcium levels and induces activation of human granulocytes. Biochem J 393, 697–704 (2006).

136. Yan, L.-J. NADH/NAD+ Redox Imbalance and Diabetic Kidney Disease. Biomolecules 11, 730 (2021).

137. Lauritzen, K. H. et al. Impaired dynamics and function of mitochondria caused by mtDNA toxicity leads to heart failure. American Journal of Physiology-Heart and Circulatory Physiology 309, H434–H449 (2015).

138. Khan, N. A. et al. mTORC1 regulates mitochondrial integrated stress response and mitochondrial myopathy progression. Cell metabolism 26, 419–428 (2017).

139. Papadopoli, D. et al. mTOR as a central regulator of lifespan and aging. F1000Res 8, F1000 Faculty Rev-998 (2019).

140. Ilter, D. et al. NADK-mediated *de novo* NADP(H) synthesis is a metabolic adaptation essential for breast cancer metastasis. Redox Biology 61, 102627 (2023).

141. Bester, A. C. et al. Nucleotide Deficiency Promotes Genomic Instability in Early Stages of Cancer Development. Cell 145, 435–446 (2011).

142. McReynolds, M. R. et al. NAD+ flux is maintained in aged mice despite lower tissue concentrations. Cell Syst 12, 1160–1172.e4 (2021).

143. McReynolds, M. R., Chellappa, K. & Baur, J. A. Age-related NAD+ decline. Exp Gerontol 134, 110888 (2020).

144. Mapuskar, K. A. et al. Mitochondrial Oxidative Metabolism: An Emerging Therapeutic Target to Improve CKD Outcomes. Biomedicines 11, 1573 (2023).

145. Console, L. et al. The Link Between the Mitochondrial Fatty Acid Oxidation Derangement and Kidney Injury. Front Physiol 11, 794 (2020).

146. Zheng Koh, D. H. & Saheki, Y. Regulation of Plasma Membrane Sterol Homeostasis by Nonvesicular Lipid Transport. Contact (Thousand Oaks) 4, 25152564211042451 (2021).

147. Tian, S., Ohta, A., Horiuchi, H. & Fukuda, R. Oxysterol-binding protein homologs mediate sterol transport from the endoplasmic reticulum to mitochondria in yeast. J Biol Chem 293, 5636–5648 (2018).

148. Tsuboi, K., Uyama, T., Okamoto, Y. & Ueda, N. Endocannabinoids and related N-acylethanolamines: biological activities and metabolism. Inflamm Regen 38, 28 (2018).

149. Boachie, N. et al. Circulating Endocannabinoids and N-Acylethanolamines in Individuals with Cannabis Use Disorder-Preliminary Findings. Brain Sci 13, 1375 (2023).

150. Peng, K.-Y. et al. Mitochondrial dysfunction-related lipid changes occur in nonalcoholic fatty liver disease progression. J Lipid Res 59, 1977–1986 (2018).

151. Schenkel, L. C. & Bakovic, M. Formation and regulation of mitochondrial membranes. Int J Cell Biol 2014, 709828 (2014).

152. Paradies, G., Paradies, V., Ruggiero, F. M. & Petrosillo, G. Role of Cardiolipin in Mitochondrial Function and Dynamics in Health and Disease: Molecular and Pharmacological Aspects. Cells 8, 728 (2019).

153. Chen, W.-W., Chao, Y.-J., Chang, W.-H., Chan, J.-F. & Hsu, Y.-H. H. Phosphatidylglycerol Incorporates into Cardiolipin to Improve Mitochondrial Activity and Inhibits Inflammation. Sci Rep 8, 4919 (2018).

154. Ahmad, T. et al. Computational classification of mitochondrial shapes reflects stress and redox state. Cell death & disease 4, (2013).

155. Long, Q. et al. Modeling of Mitochondrial Donut Formation. Biophysical journal 109, (2015).

156. Scudese, E. et al. 3D Mitochondrial Structure in Aging Human Skeletal Muscle: Insights Into MFN-2-Mediated Changes. Aging Cell **n/a**, e70054 (2025).

## References for the Supplementary Methods

157. Lam J, Katti P, Biete M, Mungai M, AshShareef S, Neikirk K, Garza-López E, Vue Z, Christensen TA, Beasley HK, et al. A Universal Approach to Analyzing Transmission Electron Microscopy with ImageJ. Cells. 2021 Aug 24;10(9):2177. doi:10.3390/cells10092177.

158. Neikirk K, Garza-López E, Marshall AG, Alghanem A, Krystofiak E, Kula B, Smith N, Shao J, Katti P, Hinton A Jr, et al. Call to action to properly utilize electron microscopy to measure organelles to monitor disease. Eur J Cell Biol. 2023 Dec;102:151365. doi:10.1016/j.ejcb.2023.151365.

159. Neikirk K, Vue Z, Katti P, Rodriguez BI, Omer S, Shao J, Christensen T, Garza-López E, Marshall A, Palavicino-Maggio C, et al. Systematic Transmission Electron Microscopy-Based Identification and 3D Reconstruction of Cellular Degradation Machinery. Adv Biol (Weinh). 2023 Jun;7(6):e2200221. doi:10.1002/adbi.202200221. (PubMed PMID: 36869426)

160. Hinton A Jr, Katti P, Christensen TA, Mungai M, Shao J, Zhang L, Trushin S, Alghanem A, Jaspersen A, Geroux RE, Neikirk K, et al. A Comprehensive Approach to Sample Preparation for Electron Microscopy and the Assessment of Mitochondrial Morphology in Tissue and Cultured Cells. Adv Biol (Weinh). 2023 Oct;7(10):e2200202. doi:10.1002/adbi.202200202. (PMCID: PMC10615857)

161. Vue Z, Neikirk K, Vang L, Garza-Lopez E, Christensen TA, Shao J, Lam J, Beasley HK, Marshall AG, Crabtree A, et al. Three-dimensional mitochondria reconstructions of murine cardiac muscle changes in size across aging. Am J Physiol Heart Circ Physiol. 2023 Nov 1;325(5):H965–H982. doi:10.1152/ajpheart.00202.2023. (Epub 2023 Aug 25)

162. Garza-Lopez, E., Vue, Z., Katti, P., Neikirk, K., Biete, M., Lam, J., Beasley, H. K., Marshall, A. G., Rodman, T. A., Christensen, T. A., Salisbury, J. L., Vang, L., Mungai, M., AshShareef, S., Murray, S. A., Shao, J., Streeter, J., Glancy, B., Pereira, R. O., Abel, E. D., … Hinton, A., Jr (2021). Protocols for Generating Surfaces and Measuring 3D Organelle Morphology Using Amira. Cells, 11(1), 65. 10.3390/cells11010065

163. Vue Z, Garza-López E, Neikirk K, Katti P, Vang L, Beasley H, Shao J, Marshall AG, Crabtree A, Murphy AC, Jenkins BC, Prasad P, Evans C, Taylor B, Mungai M, et al. 3D reconstruction of murine mitochondria reveals changes in structure during aging linked to the MICOS complex. Aging Cell. 2023 Dec;22(12):e14009. doi:10.1111/acel.14009. (Epub 2023 Nov 13)

164. Scudese E, Marshall AG, Vue Z, Exil V, Rodriguez BI, Demirci M, Vang L, Garza-López E, Neikirk K, Shao B, Le H, Stephens D, Hall DD, Rostami R, Rodman T, et al. 3D Mitochondrial Structure in Aging Human Skeletal Muscle: Insights Into MFN-2-Mediated Changes. Aging Cell. 2025 Jul;24(7):e70054. doi:10.1111/acel.70054. (Epub 2025 Apr 25; PMCID available)

